# A novel dephosphorylation peptide inhibits 17β-HSD1 enzyme activity in ovarian granulosa cells and breast cancer cells

**DOI:** 10.64898/2025.12.29.696806

**Authors:** Feng Yang, Haoyi Feng, Shanshan Chen, Xuanjun Liu, Tong Yu, Yizhao Li, Hao Wu, Shenming Zeng, Xuelei Han, Jie Lian, Kejun Wang, Xinjian Li

**Affiliations:** Department of Animal Genetics, Breeding, and Reproduction, College of Animal Science and Technology, Henan Agricultural University, Zhengzhou, China, 450046; National Engineering Laboratory for Animal Breeding, Key Laboratory of Animal Genetics and Breeding of the Ministry of Agriculture, College of Animal Science and Technology, China Agricultural University, Beijing, China, 100193; Sanya Institute of Hainan Academy of Agricultural Sciences (Hainan Laboratory Animal Research Center), Hainan, China, 571100; School of Forensic Medicine, Xinxiang Medical University, Xinxiang 453003, China; Henan Joint International Research Laboratory of Stem Cell Medicine, National Joint Engineering Laboratory of Stem Cells and Biotherapy, Xinxiang Medical University, Xinxiang 453003, China

**Keywords:** *HSD17B1*, 17β-HSD1, Enzyme activity, Phosphorylation, Breast cancer

## Abstract

Estradiol (E_2_), a pivotal mammalian reproductive hormone, is closely associated with estrogen-dependent diseases. 17β-hydroxysteroid dehydrogenase type 1 (17β-HSD1), a critical E_2_ synthesis enzyme, has been understudied for its post-translational regulation-despite its potential as a therapeutic target, no clinically approved inhibitors are currently available. In our investigation of porcine follicular atresia mechanisms, we identified five differentially phosphorylated residues in 17β-HSD1. Using in vitro site-directed mutagenesis and transgenic mouse models carrying point mutations, we established that Ser30 and Ser274 are critical phosphorylation sites that robustly modulate 17β-HSD1 enzymatic activity. We further demonstrated that activin A and insulin-like growth factor 1 (IGF-1) enhance 17β-HSD1 activity by promoting its phosphorylation at these two sites. To target these regulatory residues, we designed and synthesized cell-penetrating peptides (CPPs) that specifically suppress 17β-HSD1 phosphorylation. Functional assays conclusively demonstrated that these CPPs significantly suppressed 17β-HSD1 activity in vitro. Notably, they also inhibited the proliferation, migration and invasion of MCF-7 cells, as well as tumor growth in a mouse model of breast cancer. Our study provides novel mechanistic insights into the regulation of 17β-HSD1 and E_2_ synthesis, addressing a critical gap in steroid hormone biology. The identification of Ser30/Ser274 as functional phosphorylation sites and the development of CPP-based inhibitors offer both theoretical advances and translational potential, opening new avenues for the treatment of estrogen-dependent breast cancer.

## 1 Induction

Estradiol (E_2_) is one of the most important reproductive hormones, playing a crucial role in animal reproduction and estrogen-dependent diseases, including breast cancer ^[1]^, endometrial cancer, and ovarian cancer, endometriosis, endometrial hyperplasia, ovarian epithelial cancer, non-small cell lung cancer, and polycystic ovary syndrome ^[2–4]^. The results of “Cancer statistics, 2024” show that breast cancer is the second leading cause of cancer-associated deaths in women worldwide ^[5]^. The 17β-Hydroxysteroid dehydrogenase (17β-HSD1) is an key enzyme that converts the less active estrone (E_1_) to the more active E_2_ ^[6]^, yet its regulatory mechanism of activity remains unclear.

In the early 1950s, 17β-HSD1 was first identified in the human placenta ^[7]^. In primates, 17β-HSD1 is primarily expressed in the placenta and ovarian granulosa cells, with minor expression in the endometrium, adipose tissue, and prostate ^[8]^. 17β-HSD1 plays a crucial role in female reproductive processes and follicular development. *HSD17B1* knockout mice exhibit a normal estrous cycle and are capable of developing follicles of all stages. However, there is an increase in the ratio of E_1_/E_2_ in the ovaries, a decrease in ovarian progesterone concentrations, abnormalities in corpus luteum structure, a reduction in the number of corpus luteum, and the appearance of bundles of large granule cells of unknown origin within the ovarian matrix, all resulting in a significant decline in fertility ^[9]^. Transgenic mice overexpressing human *HSD17B1* exhibit significantly enhanced E_2_ synthesis capacity, frequently develop endometrial hyperplasia, and are unable to ovulate as adults ^[10]^. The SNP (A/C) within intron 4 of the porcine *HSD17B1* gene is associated with litter size ^[11]^. Additionally, changes in the expression level or activity of 17β-HSD1 are correlated with several estrogen-dependent diseases. 17β-HSD1 is considered a promising drug target for estrogen-dependent diseases. Inhibiting 17β-HSD1 activity can suppress the estrogen-dependent proliferation of tumor cells ^[12, 13]^. However, to date, there are no clinically available inhibitors.

Previous studies on 17β-HSD1 mostly focused on its expression and translation regulation ^[14–19]^. However, there are few studies on the regulatory mechanism of 17β-HSD1 enzymatic activity. Terhi et al. discovered that mutating His221 to Ala significantly reduces the enzyme’s oxidative (alcohol-ketone) and reductive (ketone-alcohol) activities in human 17β-HSD1, while mutating Tyr155 to Ala nearly completely suppresses the enzyme’s activity ^[20]^. The enzyme is phosphorylated at Ser134 by protein kinase A in vitro. However, phosphorylation at Ser134 does not play a role in the activity or microheterogeneity of human 17β-HSD1 ^[21]^. Research indicates that amino acid residues Ser142, Lys159, and Tyr155 are critical for the activity of 17β-HSD1 ^[21]^. Therefore, phosphorylation may affect 17β-HSD1 activity. Identifying the key phosphorylation sites of 17β-HSD1 can enhance our understanding of estrogen synthesis regulation in female animals and serve as potential drug targets for treating diseases dependent on E_2_.

During our research into the mechanisms of porcine follicular atresia, we have identified five novel sites of differential phosphorylation on 17β-HSD1 ^[22]^, whose functions have not been previously reported. We speculate that the phosphorylation of these sites may be related to enzyme activity of 17 β-HSD1, as the E_2_ level also decrease during follicular atresia. This study focused on understanding the molecular mechanisms of E_2_ synthesis regulation in ovarian follicle and breast cancer cells from the perspective of phosphorylation of the 17β-HSD1 protein.

## 2 Materials and Methods

### 2.1 Cell lines and cell culture

Primary porcine granulosa cells were isolated from ovaries of approximately 5-month-old prepubertal Large White gilts obtained from a local slaughterhouse. Immediately after excision, the ovaries were placed into a thermos flask filled with sterile normal saline pre-warmed to 37 °C and transported to the laboratory within 2 h. Upon arrival, the ovaries were washed twice with sterile normal saline containing 100 IU/mL penicillin and 50 mg/L streptomycin to remove surface debris and blood residues. Subsequently, the ovaries were immersed and gently agitated in 75% ethanol for 1 min to eliminate surface bacteria and fungi. The disinfected ovaries were then transferred into fresh sterile saline maintained at 37 °C for further processing. Healthy follicles were identified by their transparent amber appearance and vascularized follicular membrane. Follicular fluid was aspirated from visible antral follicles using a 5 mL sterile syringe. Under a stereomicroscope, cumulus-oocyte complexes (COCs) and ovarian tissues were removed. The remaining follicular fluid containing granulosa cells was centrifuged at 400 × g for 5 min at room temperature using a benchtop centrifuge. The supernatant was discarded, and the cell pellet was washed five times with PBS containing 100 IU/mL penicillin and 50 mg/L streptomycin. The final granulosa cell pellet was resuspended in DMEM/F12 (Gibco) supplemented with 10% fetal bovine serum (Gibco, 10270-106), 1% penicillin/streptomycin (Solarbio, P1400), and seeded into culture flasks. Cells were maintained in a humidified incubator at 37 °C with 5% CO₂. After 48 h, the medium was replaced to remove non-adherent cells. The adherent cells were gently washed once with PBS and cultured in fresh complete medium. Medium was subsequently changed every 24–48 h. Once a confluent monolayer was achieved, the cells were passaged for further experiments.

The HEK-293T human embryonic kidney cell line and the MCF-7 human breast cancer cell line was obtained from the American Type Culture Collection (ATCC). All cell lines were cultured in high-glucose DMEM supplemented with 10% fetal bovine serum and 1% penicillin/streptomycin at 37 °C in a humidified incubator with 5% CO₂.

### 2.2 Plasmid construction and transfection

The coding sequences of porcine and human *HSD17B1* were amplified by PCR using cDNA derived from porcine granulosa cells and human ovarian granulosa cells, respectively, as templates. The primers for amplifying *HSD17B1* and performing point mutations on it are shown in **Supplementary Table 1**. For porcine *HSD17B1*, the PCR product and pEGFP-N1 vector were digested with *EcoRI* (R3101, New England Biolabs, USA) and *BamHI* (R3136, New England Biolabs, USA). For human *HSD17B1*, digestion was performed with *EcoRI* (R3101, New England Biolabs, USA) and *AgeI-HF* (R3552, New England Biolabs, USA). The resulting fragments were ligated into the pEGFP-N1 vector to generate the recombinant plasmids pEGFP-N1-*HSD17B1* (porcine and human, respectively). The ligated constructs were transformed into *Escherichia coli* Trans1-T1 chemically competent cells (CD501-03, TransGen Biotech, China) for plasmid amplification. Plasmids were extracted using a plasmid mini-prep kit (DP103, TIANGEN Biotech, China), and the integrity and accuracy of the open reading frame (ORF) were verified by Sanger sequencing (Sangon Biotech, China). Following successful construction of the expression vectors, site-directed mutagenesis was performed to alter putative phosphorylation sites within the *HSD17B1* sequences using the Fast Site-Directed Mutagenesis Kit (FM111-01, TransGen Biotech, China), according to the manufacturer’s protocol.

Lentiviral particles were generated using a third-generation packaging system. HEK293T cells were seeded into 6-well plates and co-transfected with 3 μg of the transfer plasmid, 2.25 μg of psPAX2 (packaging plasmid), and 1 μg of pMD2.G (envelope plasmid) using Lipofectamine® 3000 (Thermo Fisher Scientific), following the manufacturer’s protocol. At 48 hours post-transfection, viral supernatants were collected and filtered through a 0.45-μm PVDF membrane to remove cellular debris. For infection, 500 μL of viral supernatant was mixed with 500 μL of fresh culture medium containing 8 μg/mL polybrene (Sigma-Aldrich) and added to target cells seeded in 12-well plates.

### 2.3 Follicle culture

The mothed of follicle culture was according to previous reports ^[23, 24]^. Ovaries from 12-to 14-day-old female mice were dissected for follicle isolation. Preantral follicles (diameter: 100–120 μm) were isolated using 33-gauge microneedles and cultured in follicle growth medium. Follicles with intact morphology were individually transferred into 0.5% alginate droplets. A pipette was used to aspirate 5–10 μL of alginate containing a single follicle. The pipette tip was held vertically 3 mm above the CaCl₂ solution surface, and gentle pressure was applied to form a spherical droplet. The droplet was touched to the CaCl₂ solution for cross-linking (2 min) to achieve solidification and encapsulation. The encapsulated follicle-alginate spheres were transferred to pre-equilibrated solution (2 mL in a 35-mm Petri dish) and equilibrated at 37°C with 5% CO₂ for 30 min. Subsequently, spheres were placed individually into wells of a 96-well plate. Each well was supplemented with 100 μL follicle growth medium (α-MEM, Solarbio, 41500) supplemented with: 1% ITS (Beyotime, C0341), 5% FBS (Gibco, A5669701), 10 mIU/mL FSH (Beijing Solarbio Science & Technology Co.,Ltd., F8470), 10 mIU/mL luteinizing hormone (Beijing Solarbio Science & Technology Co.,Ltd., L8040), and 100 U/mL penicillin-streptomycin (Solarbio, P1400). Cultures were maintained at 37°C/5% CO₂ for 6 days, with 50% medium replacement every 48 h.

Follicle images were captured on days 0, 2, and 4 using an inverted microscope. Follicle viability was assessed by morphological criteria: death was indicated by oocyte detachment from the granulosa cell layer or visible granulosa cell blackening/fragmentation. Diameters were measured using Image J via the cross-section method (two perpendicular diameters per follicle), and averages were calculated.

### 2.4 Synthetic cell-penetrating peptide (CPP) and cell treatments

The CPPs were used to specifically inhibit the phosphorylation levels of 17β-HSD1 S30 and S274 sites. The 30S CPP (RLASDPSQSF[K(Fitc)]VYATLRDLRRRRRRRR), 30A control CPP (RLASDPSQAF[K(Fitc)]VYATLRDLRRRRRRRR), 274S CPP (Rhodamine B-RMRLDDPSGSNYVTAMHRRRRRRRR) and 274A control CPP (Rhodamine B-RMRLDDPSGANYVTAMHRRRRRRRR) were synthesized by Taopu Biotech (Nanjing, China). FITC was conjugated at the K residue of 30S or 30A peptides. The Rhodamine was conjugated at N-terminal of 274S and 274A peptides. The purity of peptide was evaluated by mass spectrometry and ensured to be ≥95%. All peptides were dissolved in sterile water immediately before administration.

For pig GC and MCF-7 (estrogen-dependent breast cancer cell line) cell, different dosages of peptides were administed: 50 nM, 500 nM, 5 μM, or 50 μM. The penetrability was determined by fluorescence microscope at 0 h, 3 h, 6 h, 12 h, and 24 h after treatment (Invitrogen EVOS M5000, ThermoFisher).

Activin A were purchased from MCE (HY-P70311), 50 ng/mL activin A were used for cell treatment and follicle cultuer. IGF-1 was purchased from PrimeGene, 105-03. All other reagents were purchased from Sigma-Aldrich, unless otherwise specified.

For mouse follicle culture, 5 μM CPP was added in the culture media. The follicular diameter and survival rate were recorded every two days during the culture period.

### 2.5 Protein extraction and immunoblotting

The cell or tissue samples were harvested and washed once in PBS, then lysed on ice for 30 min with RIPA buffer (CST, 9806), and supplemented with 1% (v/v) protease inhibitor Cocktail (HY-K0010) and 1% (v/v) phosphatase inhibitors (Cocktail I, HY-K0021; Cocktail II, HY-K0022; and Cocktail III, HY-K0023), which were purchased from MCE (Shanghai, China). Western blotting was performed as described previously ^[22]^. Protein concentrations were determined using a BCA protein assay kit (Transgen Biotech, Beijing, China). Equal amounts of proteins (40-100 µg/lane) were separated by SDS-PAGE (8∼12% acrylamide running gel) and transferred to a nitrocellulose membrane (BioTrace™ NT, Pall Corp, FL, USA). The following antibodies were used in this experiment: HSD17B1 (ab134193, Abcam), Actin (ab8226, Abcam). The antibodies were diluted to the recommended ratio with Beyotime (P0256, Shanghai, China) diluent. The Western blotting images were analyzed using Image J software (National Institutes of Health, Bethesda, MD, USA). Mn^2+^-Phos-tag™ SDS-PAGE (AAL-107, FUJIFILM) was used to detect total phosphorylation of HSD17B1 according to the manufacturer’s instruction. Briefly, 5 mL of 12% (w/v) polyacrylamide: 20 μM Phos tag ™ Acrylamide gel was prepared (30% (w/v) acrylamide solution 2 mL, 1.5 mol/L Tris-HCl 1.25 mL, 5.0 mmol/L Phos-tag™ 20 μL, 10 mmol/L MnCl_2_ 20 μL, 10% (w/v) SDS 50 μL, TEMED 2 μL, 10% (w/v) ammonium persulfate solution 50 μL, and dd H_2_O 1.75 mL), and the remaining steps are similar to regular Western blotting.

### 2.6 Hormone measurement

E_2_ levels were measured using two complementary approaches. For routine quantification, E_2_ concentrations in cell culture supernatants were determined using enzyme-linked immunosorbent assay (ELISA) kits (Beijing Sino-UK Institute of Biological Technology, Beijing, China) according to the manufacturer’s instructions. The minimum detectable amount of E_2_ was 5 pg/mL, and both intra- and inter-assay coefficients of variation were <15%.

To assess the catalytic activity of wild-type and mutant 17β-HSD1, the conversion of E_1_ to E_2_ was analyzed by high-performance liquid chromatography (HPLC). HEK293T, MCF-7, and KGN cells expressing 17β-HSD1 constructs were incubated with phenol red-free DMEM/F12 medium containing 1 μg/mL E_1_ at 37°C. Supernatants were collected at defined time points, filtered through 0.22-μm PVDF membranes, and stored at –80°C. HPLC was performed using a Waters TC-C18 column (4.6 × 250 mm, 5 μm) with a mobile phase of acetonitrile: water (55:45), a flow rate of 1.0 mL/min, and UV detection at 205 nm. The conversion rate was calculated using the internal standard method based on peak area.

### 2.7 KGN cells with endogenous *Hsd17b1* knocked out

To obtain KGN cells with human *Hsd17b1* knockout, we designed gRNA-A1: ACGTCTCCAGGGATCCCGGA - GGG. The human *Hsd17b1* gene in ovarian granulosa cell line KGN was knocked out using the electroporation - mediated CRISPR/Cas9 gene editing technology. After electroporation, single clones were picked and verified by PCR and sequencing, and homozygous cells with human *Hsd17b1* gene knockout were successfully obtained. The knockout cells were identified using the upstream primer: TTGGCCGTACGTCTGGCTT and the downstream primer: TTAGGTGGGGAGACGAGGTT. Finally, KGN cells with a 58-bp deletion in *Hsd17b1* were obtained.

### 2.8 Molecular docking analysis

The molecular docking of 17β-HSD1 and its substrate E_1_ was carried out referring to the previous methods. Firstly, the protein annotation number of human 17β-HSD1, P14061, was found on the Unipt website (https://www.uniprot.org/), and the PDB file of P14061 was downloaded from the Alphafold website (https://alphafold.ebi.ac.uk/). Then, the molecular structure of E_1_ was searched using PubChem (https://pubchem.ncbi.nlm.nih.gov/). Finally, the interaction between 17β-HSD1 and E_1_ was analyzed and further visualized using Chembio3D and Autodock software.

### 2.9 Animals

All mice were housed under standard laboratory conditions with a 12 h light/dark cycle and free access to food and water. All animal procedures were approved by the Animal Care and Use Committee of Henan Agricultural University. Point-mutated *Hsd17b1* knock-in mice were generated by Jiangsu Jicui Yaokang Biotechnology Co., Ltd (Nanjing, China). Briefly, CRISPR/Cas9 components and donor vectors carrying the S30A mutation were microinjected into fertilized C57BL/6JGpt zygotes, which were then implanted into pseudo pregnant females. Genotyping of F0 offspring was performed by PCR and Sanger sequencing using primers designed against the *Hsd17b1* locus. Positive F0 founders were bred with wild-type C57BL/6JGpt mice to generate heterozygous F1 mice, followed by intercrossing to obtain homozygous (S30A^−/−^), heterozygous (S30A^+/−^), and wild-type (WT) littermates. At least five generations of backcrossing and genotyping were performed to confirm germline transmission and model stability.

To induce ovulation in mice, PMSG injections were administered at a dosage of 7.5 IU per mouse, followed by hCG injections at the same dosage 48 hours after PMSG injection. In the IGF-1 treatment group, mice were administered IGF-1 at a dosage of 2 µg/g body weight per mouse. The injections were performed both 24 hours prior to and concurrently with PMSG administration. As for the control group, mice received injections of saline (Sal) at identical time points and dosages as those used in the IGF-1 treatment for comparative analysis.

### 2.10 Cell proliferation, migration, invasion and apoptosis assay

The BeyoClick™ EdU-488 Cell Proliferation Detection Kit (C0071S, Beyotime) was employed for detecting cell proliferation. The Cell Cycle and Apoptosis Analysis Kit (C1052, Beyotime) was utilized to analyze the cell cycle. The Annexin V-FITC/PI Cell Apoptosis Detection Kit (C1062, Beyotime) was used to detect cell apoptosis. The Cell Culture Wound Closure Assay was performed following a previously described protocol ^[25]^. Briefly, for monolayer adherent cells cultured in vitro in petri dishes or plates, a micropipette tip or other hard objects were used to scratch the central area of the cell-growing region, removing the cells in the central part. Subsequently, the cells were continuously cultured until the predetermined experimental time. The transwell cell migration and invasion assay was carried out according to a previously described protocol ^[25]^, crystal violet (C0121, Beyotime) and matrigel (111010, GENOM BIO | ORG) were used for detection.

### 2.11 Tumorigenicity experiment of MCF-7 cells and treatment after tumorigenicity of CPP

Nude mice (strain, supplier, age, and sex details should be provided here) were used in this study. All mice were housed in a specific pathogen - free (SPF) environment maintained at a temperature of (25±0.5) °C and a relative humidity of (A±B) %. Water was changed every two days to ensure its freshness and prevent contamination. Mice were housed in individually ventilated cages (IVCs). Each cage was equipped with appropriate bedding material, which was changed every two days to maintain a clean and hygienic environment. The cages were placed on racks with proper air circulation to minimize the risk of disease transmission.

For the purpose of investigating the effects of CPP on breast cancer growth in nude mice, we initially employed a well-established methodology, tumorigenicity experiment of MCF-7 cells in nude mice. Firstly, referring to previous methods ^[26, 27]^, 1×10^7^ MCF-7 cells and 0.1 mL Matrix-Gel™ (Beyotime, C0372) were injected subcutaneously into nude mice to induce tumor formation. In the study, three groups were designed: the control group, the activin A treatment group, and the SS-CPP polypeptide treatment group. Initially, 10 nude mice were selected for tumorigenesis in each group. Intraperitoneal injection treatment was started when the tumor volume in nude mice reached 100∼200 mm^3^. The control group was injected with normal saline at the tumor site (n = 8). An activin A (100 ng/0.1mL/day) treatment group (n = 8), and a SS-CPP (75 µg/0.1mL/day) treatment group (n = 8). The injections were administered once a day, the size (transverse and longitudinal diameters) of the tumors was recorded every two to three days. If the diameter of a mouse’s tumor grows to more than 15 mm during the experiment, euthanize the mouse and record the size of the tumor. Finally, under the premise of meeting ethical requirements, the tumors of all mice were dissected and photographed simultaneously. The sizes of the tumor masses were compared and they were weighed. The volume of the tumor mass = Length * Width * Width * 0.5.

### 2.12 Histology

Ovarian and uterine tissues were fixed in 4% phosphate-buffered formaldehyde at 4 °C for 24 h, followed by paraffin embedding. The largest cross-sectional slices (5 μm) of the uterus and ovaries were selected for hematoxylin and eosin (H&E) staining and immunohistochemistry, as described previously. Uterine sections were analyzed for endometrial area using ImageJ software.

### 2.13 CCK8 cell viability assay

To assess the cytotoxicity of the synthetic CPPs, a CCK-8 assay (Cell Counting Kit-8, C0038, Beyotime, China) was performed according to the manufacturer’s protocol. HEK293T cells, porcine granulosa cells, and MCF-7 cells were collected, resuspended in complete culture medium, and counted using a hemocytometer. Cells were seeded into 96-well plates at a density of 3,000–5,000 cells per well in a volume of 100 μL. To prevent edge effects, PBS was added to the peripheral wells. After cell attachment, various concentrations of CPPs were added to the treatment groups. Each condition was tested in quintuplicate. Cells were incubated at 37 °C in a humidified atmosphere containing 5% CO₂ for 12, 24, or 48 h. After incubation, the medium was removed and cells were washed twice with PBS. Then, 100 μL of phenol red-free DMEM/F12 (21041025, Gibco, USA) and 10 μL of CCK-8 solution were added to each well. After an additional 60 min incubation, the absorbance at 450 nm was measured using a microplate reader. Cell viability was evaluated according to the manufacturer’s protocol.

### 2.14 Transcriptome Sequencing and Analysis

Total RNA was extracted from MCF-7 cells treated with 5 μM cell-penetrating peptide SS using TRIzol reagent (Invitrogen, USA), following the manufacturer’s instructions. RNA quality and integrity were assessed using agarose gel electrophoresis and the Agilent 5400 Bioanalyzer system (Agilent Technologies, USA). Samples with qualified integrity were used for library preparation. RNA sequencing libraries were constructed and sequenced by BIORED Biotechnology Co., Ltd. (Guangzhou, China). Library insert sizes were confirmed using the Agilent 5400 system, and the final libraries were quantified by fluorescence-based qPCR. Sequencing was performed on the Illumina NovaSeq PE150, and clean reads were obtained after filtering low-quality sequences and adapters. High-quality reads were mapped to the human reference genome GRCh38 using HISAT2, and gene expression levels were quantified and normalized as fragments per kilobase of transcript per million mapped reads (FPKM). Differential gene expression analysis was conducted using the DESeq2 R package. Genes with | log2FoldChange | > 1.5 and adjusted *p*-value < 0.05 were considered significantly differentially expressed. The raw data have been deposited in the Sequence Read Archive (SRA) database with BioProject accession number PRJNA1345717.

### 2.15 Quantitative Real-Time PCR (qRT-PCR)

To validate the transcriptome data, 1 μg of total RNA was reverse transcribed using the Evo M-MLV Reverse Transcription Premix Kit (AG11728, Accurate Biology, China), following the manufacturer’s instructions. Total RNA was extracted as described above, and reverse transcription was performed using a commercial cDNA synthesis kit (details provided in prior sections). Quantitative PCR was conducted using the SYBR Green Premix Pro Taq HS qPCR Kit (AG11701, Accurate Biology, China) according to the manufacturer’s protocol. Reactions were carried out on a CFX96 Real-Time PCR Detection System (Bio-Rad, USA). The PCR cycling conditions were: 95 °C for 30 s, followed by 40 cycles of 95 °C for 5 s and 60 °C for 15 s. Each sample was analyzed in triplicate. The housekeeping gene GAPDH was used for normalization. Relative gene expression levels were calculated using the 2^^−ΔΔCt^ method.

### 2.16 Statistical analysis

All statistical analyses were performed using SPSS 22.0 software (IBM, USA). Data are presented as mean ± SD. Differences (the gray values of protein expression, hormone levels, tumor size, tumor weight, body weight of mice, etc.) between groups were assessed using ANOVA, followed by an LSD post hoc test. The t - test was used to analyze the differences between the two groups in terms of indicators such as uterine area, follicle diameter, gray values of some proteins, E_2_ levels, and relative mRNA expression. *P*< 0.05 was considered statistically significant.

## 3 Results

### 3.1. 17β-HSD1 phosphorylation level decreased during porcine follicular atresia

Our previous proteomic and phosphorytomic studies revealed a significant reduction in the phosphorylation level of five sites (Ser30, Ser269, Ser274, Ser275, and Thr319) of 17β-HSD1 in granulosa cells as porcine follicles transitioned from healthy to atretic; however, there was no change in the protein level of 17β-HSD1 ^[22]^ (**Fig. 1A and 1B**). Based on the higher phosphorylation levels of 17β-HSD1 at these five sites in healthy follicles, we hypothesized that the overall phosphorylation level of 17β-HSD1 is also elevated in healthy follicles. To investigate this hypothesis, we utilized Phos-tag SDS-PAGE to assess the total phosphorylation level of 17β-HSD1. As anticipated, our results demonstrated that the protein expression level of 17β-HSD1 was comparable in granulosa cells from healthy, slightly atretic, and atretic follicles; however, the total phosphorylation level of 17β-HSD1 in granulosa cells from healthy follicles was significantly higher than that observed in slightly atretic and atretic follicles (**Fig.1C**). Among the five sites Thr319 exhibited the largest change ratio (H/A ratio =4.144), while Ser30 and Ser274 are conserved sites in humans, pigs, and mice (**Fig. S1A**). Consequently, these three specific antibodies were customized for these three sites to verify their respective phosphorylation levels. The results showed that the phosphorylation levels at these three sites were indeed consistent with the results of our team’s previous phosphorytomics (**Fig. 1D**).

**Fig. 1.**
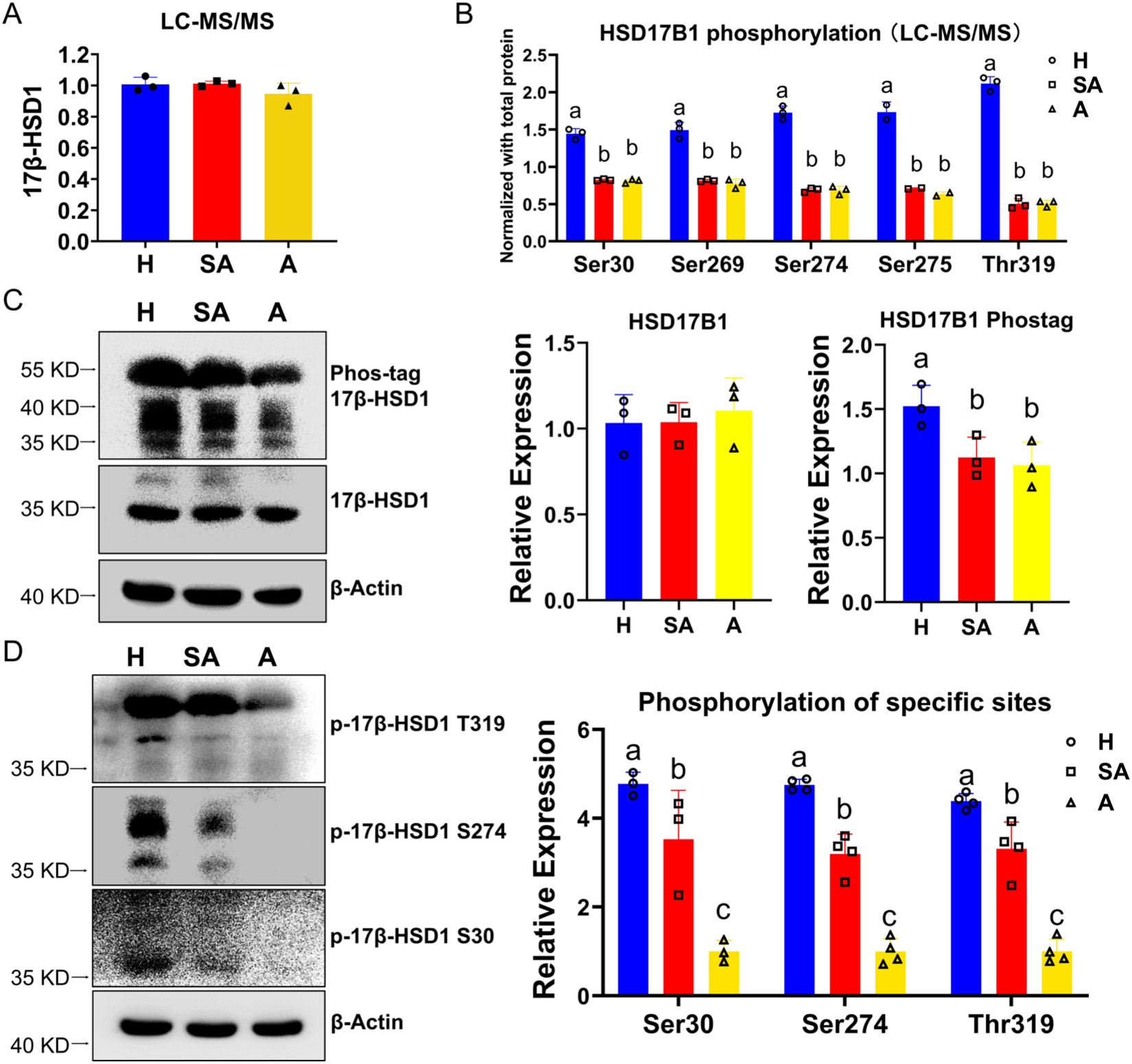
Validation of novel phosphorylation site changes in 17β-HSD1 during follicular atresia A. Changes in 17β-HSD1 detected in proteomics during the process of porcine follicular atresia within granulosa cells. B. Changes in phosphorylation levels at five sites of 17β-HSD1 detected in phosphoproteomics during the process of porcine follicular atresia. C. Detection of protein expression and total phosphorylation changes of 17β-HSD1 in granulosa cells during porcine follicular atresia using immunoblotting. For the detection of total phosphorylation, gels with added Phos-tag were utilized. D. Detection of phosphorylation changes at three sites of 17β-HSD1 during porcine follicular atresia using customized phosphorylated antibodies. The experiment was repeated at least three times. The different superscript letters in the bar graph indicate significant differences between groups. H= healthy porcine follicle; SA= slightly atretic follicle; A= atretic follicle.

### 3.2 Dephosphorylation of 17β-HSD1 at Ser 30 and Ser 274 depressed its enzyme activity

We speculate that dephosphorylation of 17β-HSD1 may depress its enzyme activity, as the E_2_ level also decrease during follicular atresia. To investigate the potential impact of these five sites on the enzymatic activity of 17β-HSD1, we constructed expression vectors harboring mutations at these sites. Specifically, Ser or Thr residues were mutated to Ala to prevent phosphorylation at these sites, and to Glu to mimic phosphorylation. To determine if the corresponding sites in human 17β-HSD1 also influence enzymatic activity, we aligned the amino acid sequences of 17β-HSD1 from human, pig, and mouse, revealing that Thr319 is a unique phosphorylation site in pigs, while the analogous sites in humans are Ser30 and Ser274, as shown in **Fig. S1A**. Consequently, we also generated expression vectors with Ser30 and Ser274 mutations in human 17β-HSD1, with schematic representations presented in **Fig. S1B and C**. Western blotting analysis confirmed the successful expression of fusion proteins following transfection of these vectors into 293T cells, as depicted in **Fig. 2A and B**. Subsequently, we evaluated the enzymatic activity of 17β-HSD1 by transfecting cells with vectors harboring mutations at various sites, adding 1 ng/mL E_1_ as a substrate to the cell culture medium, and measuring E_2_ production at different time points post-addition, as shown in **Fig. 2C**. Our results indicate that at 0.5 hours, the E_2_ concentrations in the dephosphorylated Ser30 and Ser274 mutants, as well as the empty vector group, were significantly lower than those in the wild-type 17β-HSD1 group. Moreover, compared to the phosphorylated mimics of Ser30, Ser274, and Thr319, dephosphorylation at these sites also significantly inhibited E_2_ synthesis (**Fig. 2D**). These findings suggest that phosphorylation of Ser30, Ser274, and Thr319 is essential for maintaining the enzymatic activity of 17β-HSD1. At 1.5 hours post-medium change, however, no significant differences were observed among the various mutant groups, likely due to the exhaustion of the substrate E_1_ in the culture medium. To more rigorously examine the impact of site mutations on enzyme activity, we transfected KGN cells with endogenous *Hsd17b1* knocked out using lentiviral vectors (**Fig. S2**) and employed high-performance liquid chromatography (HPLC) to detect the synthesis of E_2_. Results also showed that dephosphorylation of pig 17β-HSD1 at Ser30 and Ser274 can inhibit its enzyme activity, and mimetic phosphorylation mutations at the corresponding sites can partially restore the enzymatic activity of 17β-HSD1, **Fig 2. E, F**.

**Fig. 2.**
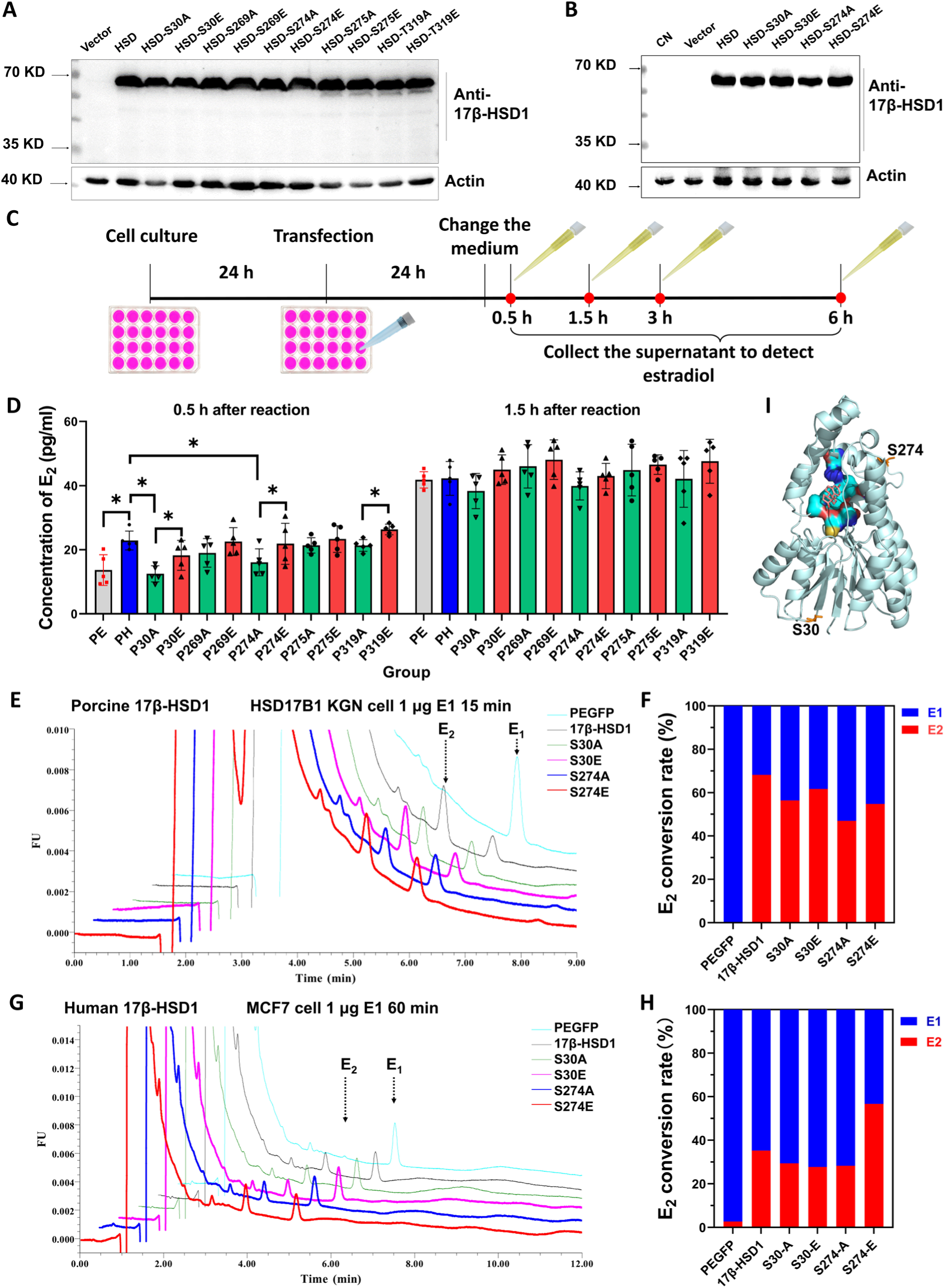
Ser30 and Ser274 are key phosphorylation sites that regulating 17β-HSD1 activity A. After transfection of 293T cells with the pEGFP-N1 empty vector, porcine 17β-HSD1 and its mutant vectors for 48 hours, the protein expression of 17β-HSD1 was detected. B. After transfection of 293T cells with the pEGFP-N1 empty vector, human 17β-HSD1 and its mutant vectors for 48 hours, the protein expression of 17β-HSD1 was detected. C. The experimental flowchart of transfecting 293T cells with porcine 17β-HSD1 and its different mutant vectors and detecting the enzymatic activity of 17β-HSD1. After plasmid transfection for 24 hours, the medium was replaced with phenol red-free medium containing 1 ng/mL E1, and then the supernatant was taken at different times to detect the synthesis of E2, that is, the enzymatic activity of 17β-HSD1. D. The concentration changes of E2 in the culture medium at 0.5 hours and 1.5 hours after the addition of E1. The concentration of E2 in the culture medium was detected by ELISA kit. E. Porcine 17β-HSD1 and its mutant vectors were transfected into HSD17B1-knockout KGN cells. After 48 hours of transfection, a phenol red-free medium containing 1 ng/mL E1 was added, and the E2 conversion rate was detected after a 15-minute reaction. F. Quantitatively show the E2 conversion rate after porcine 17β-HSD1 and its mutations at different sites. G. Human 17β-HSD1 and its mutant vectors were transfected into MCF7 cells. After 48 hours of transfection, a phenol red-free medium containing 1 ng/mL E1 was added, and the E2 conversion rate was detected after a 60-minute reaction. H. Quantitatively show the E2 conversion rate after human 17β-HSD1 and its mutations at different sites. I. The molecular docking diagram of E1 and human 17β-HSD1, in which the positions of the two amino acid residues, Ser30 and Ser274, were marked.

However, after transfecting human 17β-HSD1 and its mutant vectors into 293T cells not significantly influence E_2_ conversion rate (**Fig. S3**), which might be caused by the inappropriate cell type. Therefore, we transfected human 17β-HSD1 and its mutant vectors into MCF-7 cells, an estrogen-dependent human breast cancer cell line, and detected the enzyme activity by HPLC ^[26]^. The results showed that after adding E_1_ for 60 minutes, the mimetic phosphorylation mutation at the Ser274 site could increase the E_2_ conversion rate by 21.4%, but the mutation at the Ser30 site did not show an obvious change (**Fig 2. G, H**). This indicates that Ser274 has a strong regulatory ability on the enzyme activity of human 17β-HSD1, while Ser30 has a weak regulatory ability on the enzyme activity of human 17β-HSD1.

Therefore, Ser30 and Ser274 might be the key phosphorylation sites for regulating the enzymatic activity of 17β-HSD1, but whether their mutations affect the binding of 17β-HSD1 to the substrate E_1_ is unknown. We found through molecular docking that Ser30 and Ser274 are not on the binding pocket of 17β-HSD1 to the substrate E_1_, **Fig 2. I**. Therefore, mutation at these two sites do not directly affect the binding of 17β-HSD1 to E_1_, and it is scientifically reliable to study whether the phosphorylation of these two sites regulates its enzymatic activity through site mutation.

### 3.3 Construct 17β-HSD1 S30A site mutant mice and detect their phenotypes

Furthermore, to delve deeper into the functional implications of phosphorylation at the Ser30 site, we employed CRISPR-CAS9 technology to generate a point-mutant mouse model harboring a 17β-HSD1 S30A mutation (**Fig. 3A**). During the inbreeding process of point-mutant mice, we found that the litter size of S30A homozygous mutant mice was significantly lower than that of heterozygous and wild-type mice, as shown in **Fig. 3 B**. We found that there was no significant difference in the expression of 17β-HSD1 between S30A homozygous mutant mice and wild-type mice, while the phosphorylation level of 17β-HSD1 Ser30 was significantly reduced in the uterus, which further proved the success of the point mutation, as shown in **Fig. 3 C, D**. Examination of ovarian and uterine sections of point-mutant mice revealed that the cross-sectional area of the uterus in mutant mice was reduced compared with the control group, as shown in **Fig. 3 E, F**. Subsequent analysis of serum hormones revealed that while the E_2_/E_1_ ratio did not decrease significantly, the serum E_2_/P ratio was markedly reduced in point-mutated mice (**Fig. 3G**). Mice with point mutations can come into heat, but the estrous cycle is prolonged or irregular. These observations underscore the pivotal role of Ser30 phosphorylation in modulating 17β-HSD1 activity in vivo, as well as its subsequent impact on estrogen homeostasis and reproductive function.

**Fig. 3.**
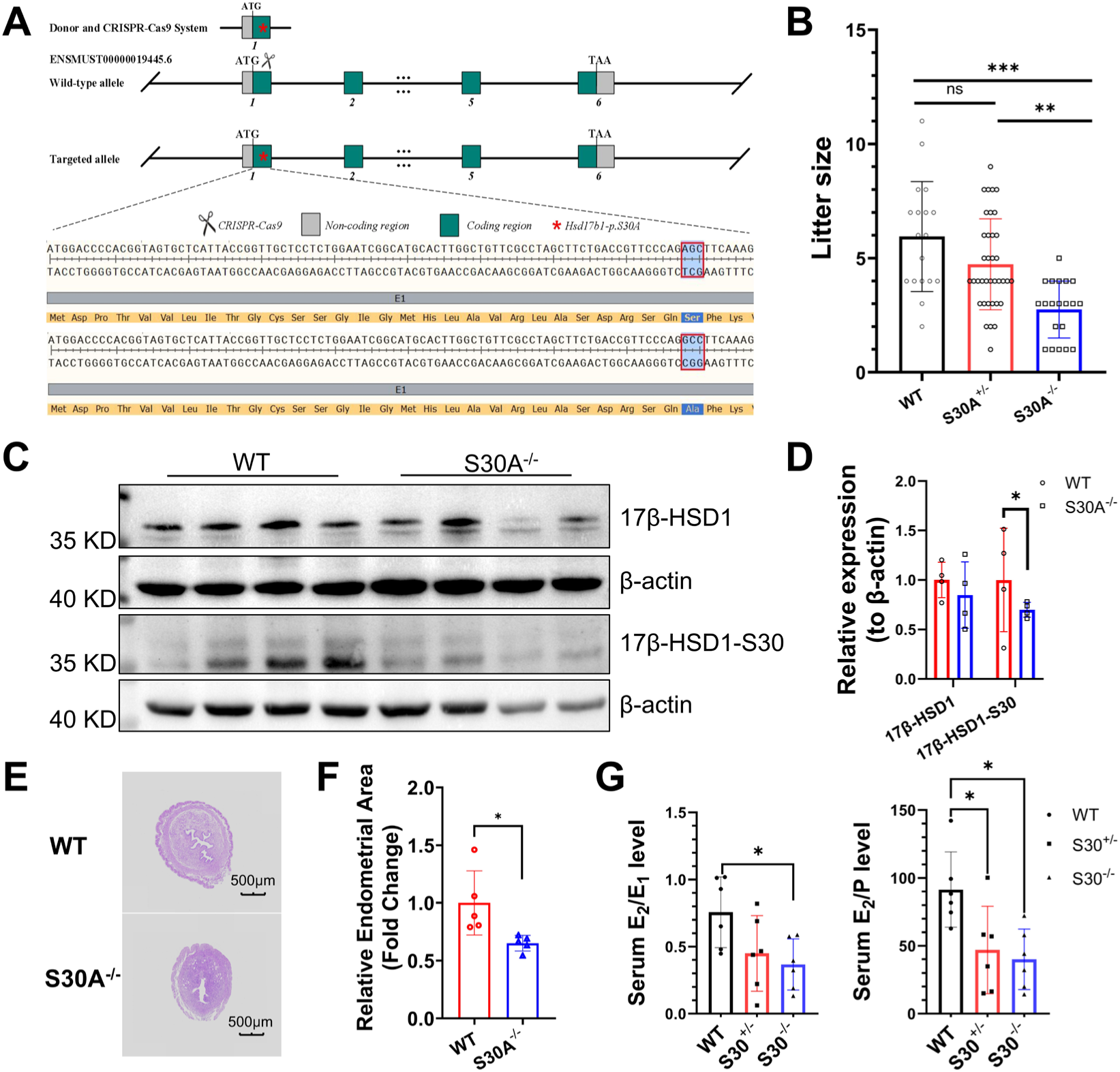
The 17β-HSD1 S30A site mutant mice show decrease of E2 level and offspring number A. Schematic diagram of constructing 17β-HSD1 S30A mutation mice using CRISPR-CAS9 technology. B. Comparison of litter sizes among wild-type, heterozygous, and homozygous mice. C, D. Comparison of 17β-HSD1 expression and 17β-HSD1 Ser30 phosphorylation levels in the uterus of wild-type and homozygous mice. E, F. Comparison of uterine sections and cross-sectional areas between wild-type and homozygous mice. G. Comparison of serum E2 levels between wild-type and homozygous mice. The “*” indicates a significant difference between groups.

### 3.4 Activin A and IGF-1 promoted 17β-HSD1 phosphorylation and its enzymatic activity

The synthesis of E_2_ in granulosa cells is dependent on pituitary gonadotropins and local ovarian signaling molecules like activin A and insulin-like growth factor 1 (IGF-1) ^[28, 29]^. Although activin has been reported to exert a positive effect on 17β-HSD1 activity ^[28]^, the underlying mechanism remains incompletely understood. Mice with knockout of activin subunits are infertile ^[30]^. Recent studies have demonstrated that a decrease in the ratio of activin to inhibin can induce apoptosis in porcine granulosa cells, leading to the promotion of follicular atresia ^[31, 32]^. In human, activin A is regarded as a cancer modulator that is involved in carcinogenesis in various tissues and organs ^[33]^, such as ovarian cancer ^[34]^ and breast cancer ^[35–37]^. Whether activin A and IGF-1, via their regulation of 17β-HSD1 phosphorylation, can affect its enzymatic activity, thereby modulating E_2_ synthesis and ultimately influencing follicular development and the cancer process remains a fascinating question.

By adding 50 ng/mL of activin A in the cultured porcine ovarian granulosa cells and detecting the expression and phosphorylation changes of 17β-HSD1, the results indicated that activin A treatment has no significant effect on the expression of 17β-HSD1, but upregulated the phosphorylation level of 17β-HSD1 at Ser30, Ser274, Thr319, and total phosphorylation level of 17β-HSD1, as shown in **Fig. 4 A-F**. Moreover, activin A treatment significantly increased the synthesis of E_2_ in the culture medium of porcine ovarian granulosa cells, as shown in **Fig. 4 G**. Similarly, treatment of MCF-7 cells with 50 ng/mL activin A also significantly increased the phosphorylation levels of 17β-HSD1 at Ser30 and Ser274, as well as the total phosphorylation level of 17β-HSD1, as shown in **Fig. 4 H, I**. Furthermore, when we added 50 ng/mL activin A during the three-dimensional culture of mouse follicles, we found that activin A could improve the survival rate and growth of in vitro cultured follicles, as shown in **Fig. 4 J-L**.

**Fig. 4.**
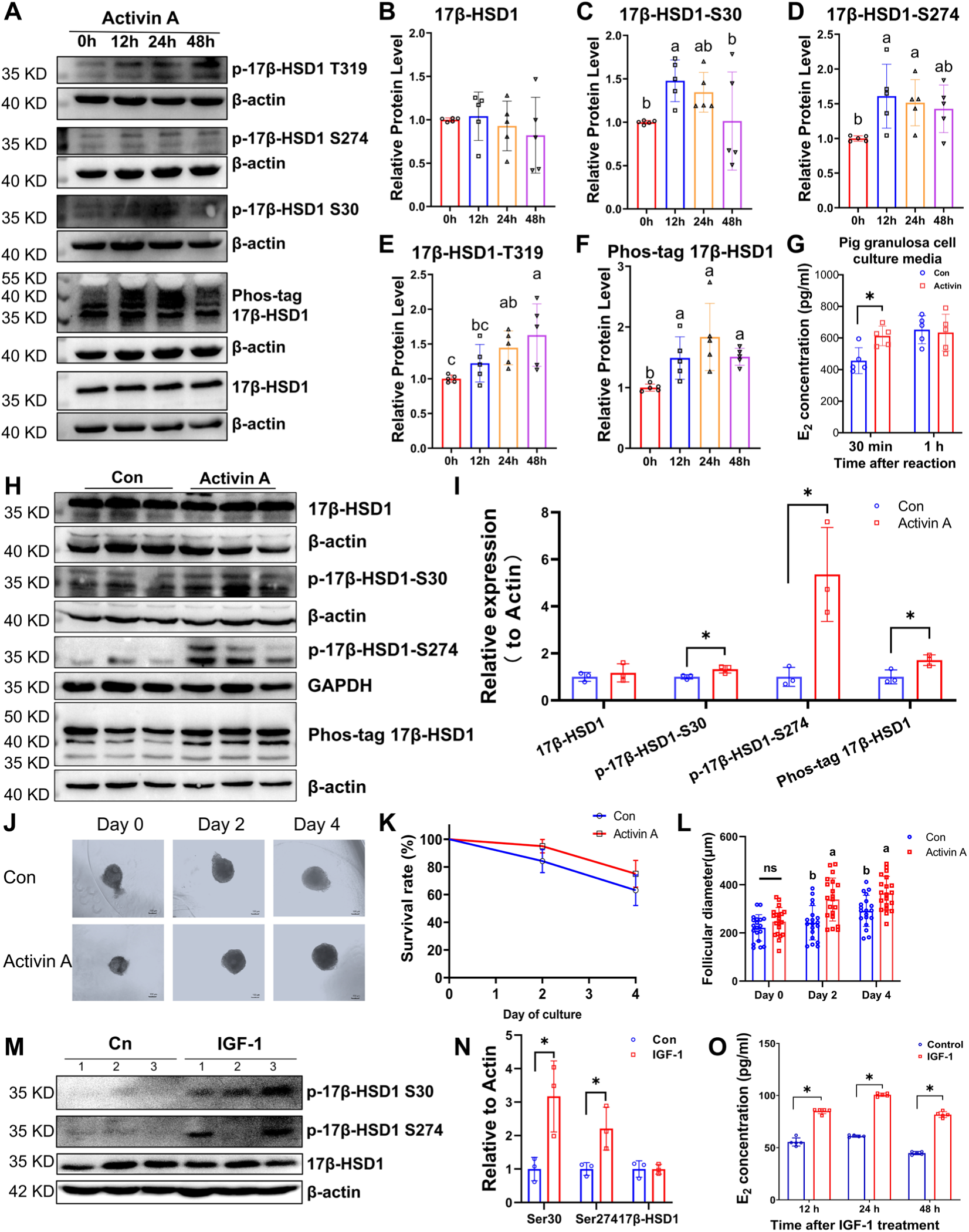
Activin A and IGF-1 can facilitate 17β-HSD1 phosphorylation and E2 synthesis A. After treating porcine granulosa cells with activin A for different time periods, Western blot was used to detect the expression of 17β-HSD1, phosphorylation at key sites, and the total phosphorylation level. The experiment was repeated five times. B-F. Gray value analysis of the expression of 17β-HSD1, phosphorylation at key sites, and the total phosphorylation level. Different superscript letters indicate significant differences between groups. G. Changes in the concentration of E2 in the cell culture medium of porcine granulosa cells after treating with activin A for 24 h. H. Detection of the protein expression of 17β-HSD1, 17β-HSD1 Ser30, 17β-HSD1 Ser274, and total phosphorylation of 17β-HSD1 in MCF-7 cells treated with activin A (50 ng/mL) using Western blotting; I. Analysis of the relative expression levels of 17β-HSD1 and its phosphorylation by densitometry; J. Representative pictures of three-dimensional culture of mouse follicles, bar=100 μm. K. Comparison of follicle survival rates after activin A treatment. L. Comparison of follicle growth diameters after activin A treatment. M. After injecting 80 μg IGF-1 into KM mice for 24 hours, Western blot was used to detect the expression of 17β-HSD1 and the phosphorylation level at key sites in the ovary. N. Quantitative analysis of the gray value in M. O. Blood was collected at different time points after injecting 80 μg IGF-1 into KM mice, and the concentration of serum E2 was detected. Five mice were repeated at each time point. The “*” indicates a significant difference between groups.

Additionally, after intraperitoneal injection of 80 μg of IGF-1 into KM mice, there is no significant difference in the expression of ovarian 17β-HSD1, but the phosphorylation levels at Ser30 and Ser274 sites are significantly increased, as shown in **Fig. 4 M**, **N**, and the concentration of E_2_ in the mouse serum is significantly increased after IGF-1 treatment, as shown in **Fig. 4 O**.

### 3.5. Cell penetrating peptide (CPP) treatment hindered 17β-HSD1 enzyme activity

Due to the crucial role that E_2_ plays in the development and progression of estrogen-dependent diseases such as hormone-dependent breast cancer and ovarian cancer, 17β-HSD1 is thus regarded as an important drug target for the treatment of these diseases ^[38]^. However, as of now, there are no clinically available 17β-HSD1 inhibitors.

Cell-penetrating peptides (CPPs) are oligopeptides consisting of approximately 8∼30 amino acid residues and have the ability to be taken up by living cells ^[39]^. CPPs can transport various biologically active substances into cells, including proteins, peptides, DNA, siRNA, and small molecule drugs, thereby enabling these carried substances to exert their functions. CPPs have been applied to inhibit the phosphorylation of proteins to treat certain diseases. For example, CPPs have been used to inhibit hyperphosphorylated tau, providing a new idea for the targeted treatment of Alzheimer’s disease ^[40]^. Therefore, we used synthetic peptide Ser30-R8 (30S CPP) or Ser274-R8 (274S CPP), which contains the unique Ser30 or Ser274 phosphorylation site of 17β-HSD1in its C terminus and a cell-penetrating sequence (R8) in the N terminus to allow for cellular internalization of the peptide. This peptide can inhibit the phosphorylation of 17β-HSD1 at Ser30 or Ser274 by competitive binding with its kinase. The Ser30A-R8 (30A CPP) or Ser274A-R8 (274A CPP) peptide, in which Ser30 or Ser274 is mutated to Ala, was used as control, and the sequence is shown in **Fig. 5 A**. Among them, Ser30 was labeled with FITC, and Ser74 was labeled with Rhodamine B. The molecular structural formula of the transmembrane polypeptide is shown in **Fig. S4**. Firstly, we detected the cell-penetrating ability of transmembrane polypeptides at different concentrations (**Fig. S5, S6**), and the results showed that the 5 μM transmembrane polypeptide could effectively enter porcine granulosa cells and MCF-7 cells, **Fig. 5 B**. Next, we detected the changes in cell viability (CCK-8) after treating porcine granulosa cells, MCF-7 cells, and 293T cells with 5 μM transmembrane polypeptides for different times, and found that the treatment with the transmembrane polypeptide for 12 h and 24 h could inhibit the viability of porcine granulosa cells and MCF-7 cells, and the inhibitory ability weakened after 48 h of treatment. However, the transmembrane polypeptide had no inhibitory effect on the viability of 293T cells, **Fig. 5 C**. This indicates that transmembrane polypeptides exert cell-specific effects on cell viability and tend to inhibit cells with estrogen-dependent proliferation. After treating porcine granulosa cells with the transmembrane polypeptide for 3 h and 12 h, the E_2_ synthesis ability of the cells was significantly inhibited after the addition of E_1_ for a reaction of 0.5 h or 1 h. Among them, the combined use of the transmembrane polypeptides at the two sites, which is, the SS group, resulted in the greatest reduction in E_2_ synthesis compared with the control group, **Fig. 5 D, E**. After 3 hours of CPP treatment, the activity of 17β-HSD1 enzyme can be inhibited, indicating that CPPs has a highly efficient transmembrane ability, and the inhibitory effect of CPPs on the activity of 17β-HSD1 enzyme is not caused by reducing cell activity. To further detect the effect of the SS transmembrane polypeptide treatment on the enzymatic activity of MCF-7, we first overexpressed human 17β-HSD1 in MCF-7 cells, and then treated the cells with or without the SS transmembrane polypeptide, and then added the substrate E_1_, and detected the E_2_ conversion rate by the HLPC method. The HLPC results were shown in **Fig. 5 F, G**, and the statistical results indicate that the SS group transmembrane polypeptide treatment significantly reduced the E_2_ conversion rate of MCF-7. The results demonstrated that the treatment with target CPPs can significantly inhibit the 17β-HSD1 enzyme activity in both porcine granulosa cells and MCF-7 cells.

**Fig. 5.**
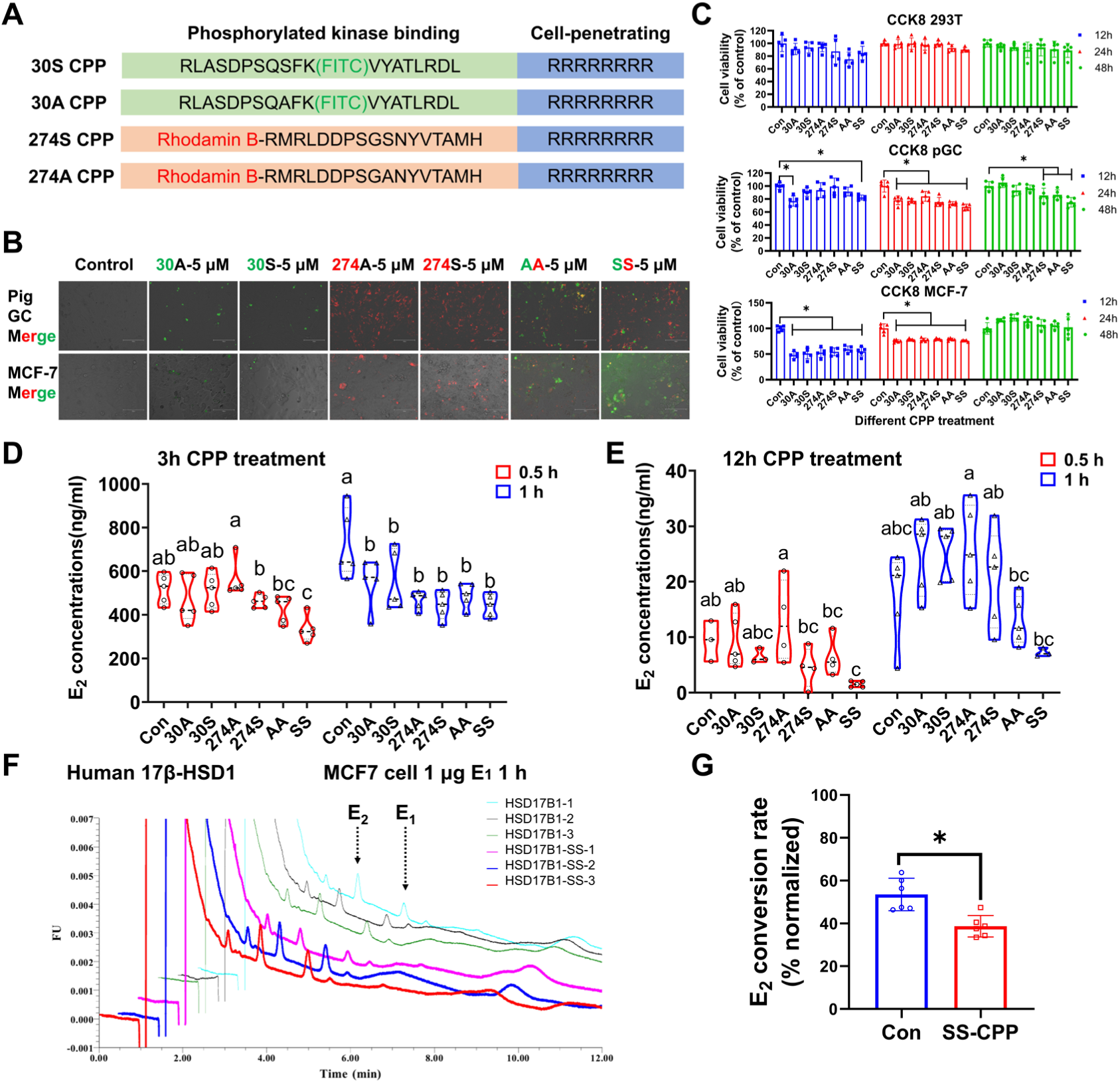
CPP treatment hindered 17β-HSD1enzyme activity A. Transmembrane polypeptides designed for Ser30 and Ser74 of 17β-HSD1. FITC represents fluorescein isothiocyanate. For fluorescein labeling, FITC is combined with the K18 residue of the polypeptide; Rhodamine B represents an artificially synthesized fluorescent dye, which is connected to the N-terminal of the polypeptide. R8 is the transmembrane sequence. B. After treating porcine granulosa cells and MCF-7 cells with 5 μM CPP for 24 hours, the situation of CPP entering the cells is detected by fluorescence microscopy. Bar=150 μm. C. The cell viability changes of porcine granulosa cells and MCF-7 cells treated with 5 μM CPP for different times are detected by CCK8. D, E. After treating the cultured porcine granulosa cells with 5 μM CPP for 3 h or 12 h, the synthesis of E2 in the cell culture medium is detected. F. After MCF-7 cells transfected with human 17β-HSD1 for 24 hours, two CPPs are added for combined treatment (SS group) or no CPP is added (control group) for 12 hours, and then 1 μg of the substrate E1 is added. The conversion rate of E2 is detected by HLPC. G. The difference in the conversion rate of E2 between the transmembrane peptide treatment in the SS group and the control group is statistically analyzed. Different superscript letters indicate significant differences between groups. The “*” indicates a significant difference between groups.

### 3.6. CPP treatment inhibited activin A induced 17β-HSD1 phosphorylation

In order to verify whether CPP affects the enzymatic activity of 17β-HSD1 by inhibiting its phosphorylation, we added different CPP treatments after treating porcine granulosa cells with activin A, and detected the phosphorylation of Ser30 and Ser274 sites of 17β-HSD1 and the changes in its total phosphorylation by western blotting, **Fig. 6 A**. The results showed that the phosphorylation level of the Ser30 site after CPP treatment in the 30S group did not decrease significantly, while the phosphorylation level of the Ser30 site after CPP treatment in the SS group decreased significantly, **Fig. 6 B**. Compared with the activin A treatment group, both the 274S and 274A CPP treatment groups could significantly inhibit the phosphorylation level of the Ser274 site, and the phosphorylation level of the Ser274 site after CPP treatment in the SS group decreased more significantly, as shown in **Fig. 6 C**. After CPP treatment in the AA group and the SS group, the total phosphorylation level of 17β-HSD1 decreased significantly, as shown in **Fig. 6 D**.

**Fig. 6.**
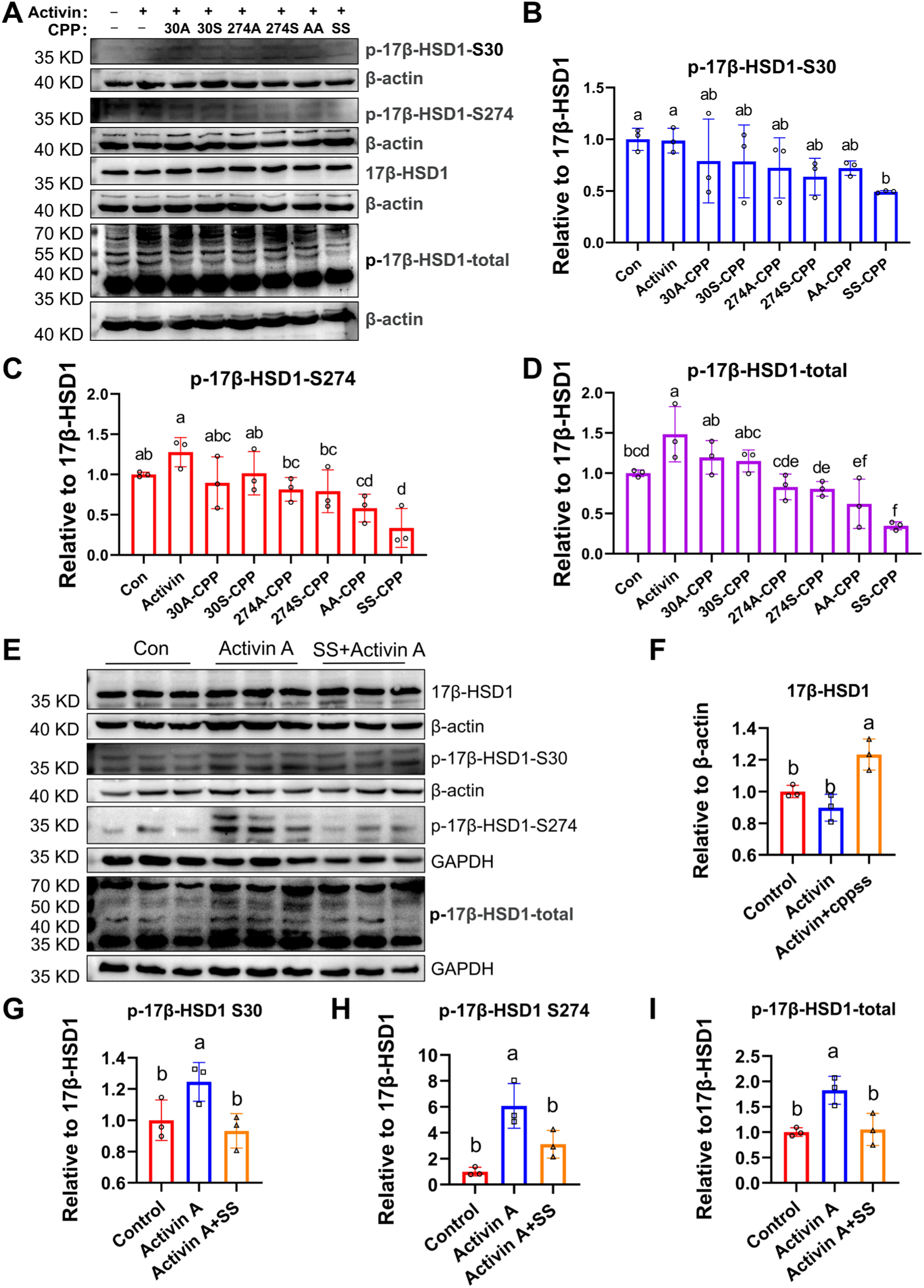
The treatment of SS-CPP reduces the phosphorylation level of 17β-HSD1 A. The effects of porcine granulosa cells treated with and without activin A and different CPPs on the phosphorylation at the key sites and total phosphorylation of 17β-HSD1 were investigated. B. The relative quantitative analysis of the expression of 17β-HSD1 Ser30 site relative to 17β-HSD1. C. The relative quantitative analysis of the expression of 17β-HSD1 Ser274 site relative to 17β-HSD1. D. The relative quantitative analysis of the total phosphorylation level of 17β-HSD1 relative to 17β-HSD1 expression. E. Detection of changes in 17β-HSD1 and its phosphorylation levels in MCF-7 cells after co-treatment with cell-penetrating peptides and activin A for 12 hours using Western blotting. F. Analysis of relative expression levels of 17β-HSD1 by densitometry. G. Analysis of phosphorylation levels of 17β-HSD1 at Ser30 relative to 17β-HSD1 expression by densitometry. H. Analysis of phosphorylation levels of 17β-HSD1 at Ser274 relative to 17β-HSD1 expression by densitometry. I. Analysis of total phosphorylation levels of 17β-HSD1 relative to 17β-HSD1 expression by densitometry. Different superscript letters indicate significant differences between groups.

Subsequently, we verified whether CPPs can inhibit the phosphorylation of 17β-HSD1 in MCF-7 cells, the results were shown in **Fig. 6 E-I**. The SS-CPP combination treatment significantly increased 17β-HSD1 protein expression in MCF-7 cells (*P* < 0.05), while suppressed activin A-induced Ser30, Ser274, and total 17β-HSD1 phosphorylation levels (*P* < 0.05). This result indicated that SS-CPP can inhibit the enzymatic activity of 17β-HSD1 by suppressing the phosphorylation level of 17β-HSD1 at the key sites.

### 3.7. SS-CPP inhibited the proliferation, migration and invasion of MCF-7 cell

To further investigate whether SS-CPP has the potential to inhibit the progression of breast cancer, we examined the changes in gene transcription, cell proliferation, apoptosis, migration and invasion after treating MCF7 cells with SS CPP.

The transcriptome results showed that there were 1321 differentially expressed genes, including 575 up-regulated genes and 746 down-regulated genes. The genes altered by SS CPP treatment were significantly enriched in pathways related to cell proliferation, migration, apoptosis, and cancer (*P* < 0.05), as shown in **Fig. S7 A-C**. Moreover, qPCR was used to detect the expression of genes related to proliferation and apoptosis in MCF-7 cells treated with the SS CPP group, as shown in **Fig. S7 D**. The results confirmed that the SS CPP group inhibited the proliferation of MCF - 7 cells and verified the reliability of the sequencing data.

### 3.8. SS-CPP inhibited the growth of tumors in MCF-7 breast cancer-bearing nude mice

To further investigate whether SS-CPP has the potential to inhibit the progression of breast cancer, we examined the growth changes of cancer tissues after tumor formation in nude mice.

In the nude mouse tumorigenicity experiment, initially, 10 nude mice were used for tumorigenesis in the control group, the activin A treatment group, and the SS-CPP treatment group. Finally, 8 nude mice in each group successfully formed tumors. Drug treatment was started on the 21st day after tumor formation. Two mice in the control group were terminated from the experiment on the 30th day after tumor formation because the tumor diameter exceeded 15 mm. In the activin A treatment group, two mice were terminated on the 28th day and one mouse was terminated on the 30th day after tumor formation due to the tumor diameter exceeding 15 mm. In the SS-CPP treatment group, no mice had a tumor diameter exceeding 15 mm during the experiment, but the tumor of one mouse regressed midway. Therefore, when the tumors were dissected on the 31st day, there were 6 mice in the control group, 5 mice in the activin A treatment group, and 7 mice in the SS-CPP treatment group, as shown in **Fig. 8A**. After dissecting the tumors on the 31st day and weighing them, it was found that the tumor weight in the SS-CPP treatment group was significantly lower than that in the activin A treatment group, and the average tumor weight was also lower than that in the control group, as shown in **Fig. 8B**. From the plotted tumor growth curve, it can also be seen that the volume, length, and width of the tumors in the SS-CPP treatment group were smaller than those in the activin A treatment group and the control group, as shown in **Fig. 8 C-E**. This indicates that SS-CPP treatment can inhibit the growth of tumors formed by MCF-7. There was no significant difference in the body weight changes of mice in the SS-CPP treatment group compared with those in the activin A treatment group and the control group, as shown in **Fig. 8 F**. This sheds new light on the treatment of breast cancer.

**Fig. 7.**
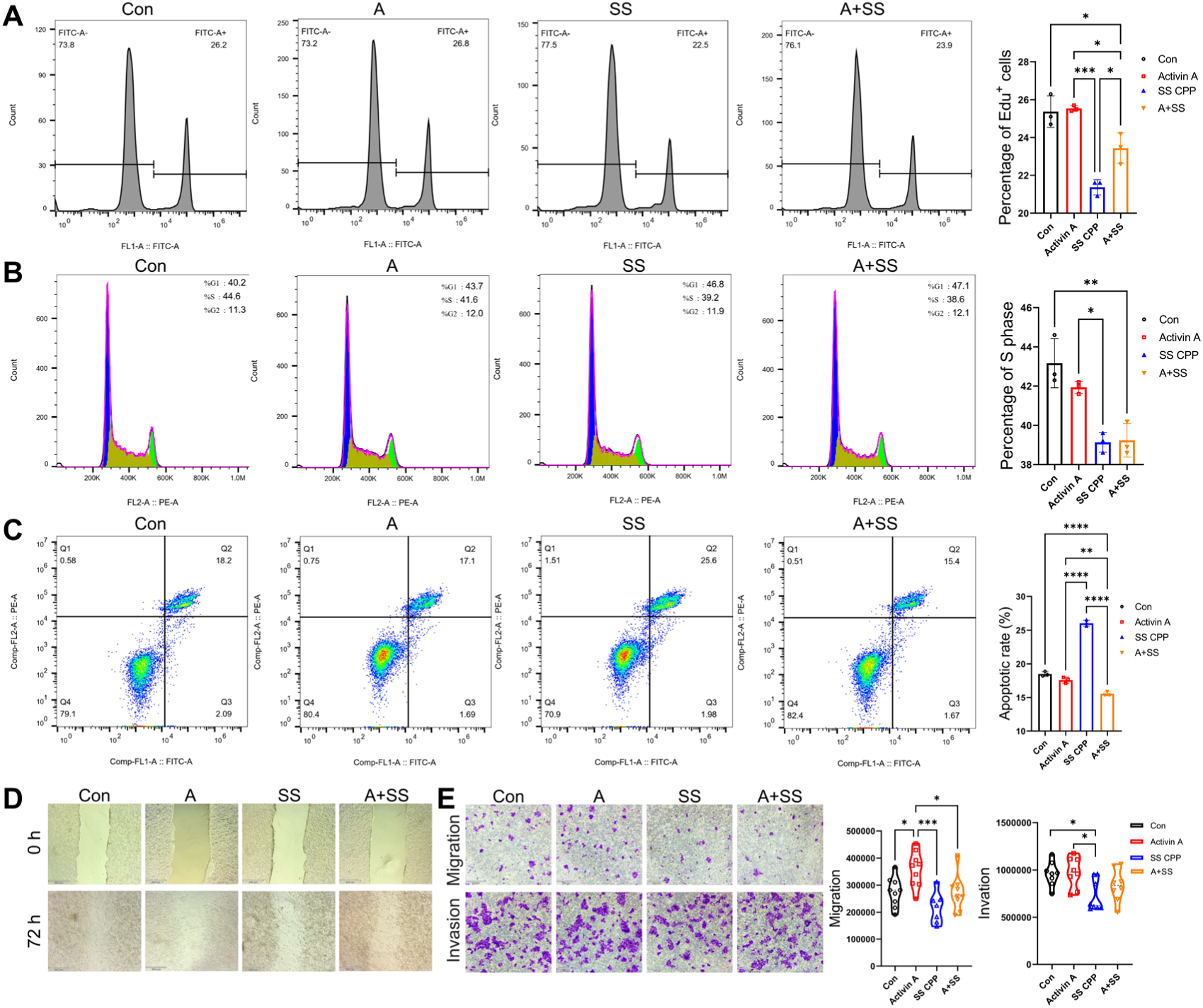
Effect of SS-CPP treatment on the proliferation, apoptosis, migration and invasion of MCF 7 cell A. Effect of SS-CPP treatment on the proliferation of MCF 7 cell. B. Effect of SS-CPP treatment on the cell cycle of MCF 7 cell. C. Effect of SS-CPP treatment on the apoptosis of MCF 7 cell. D. Effect of SS-CPP treatment on the proliferation of MCF 7 cell.

**Fig. 8.**
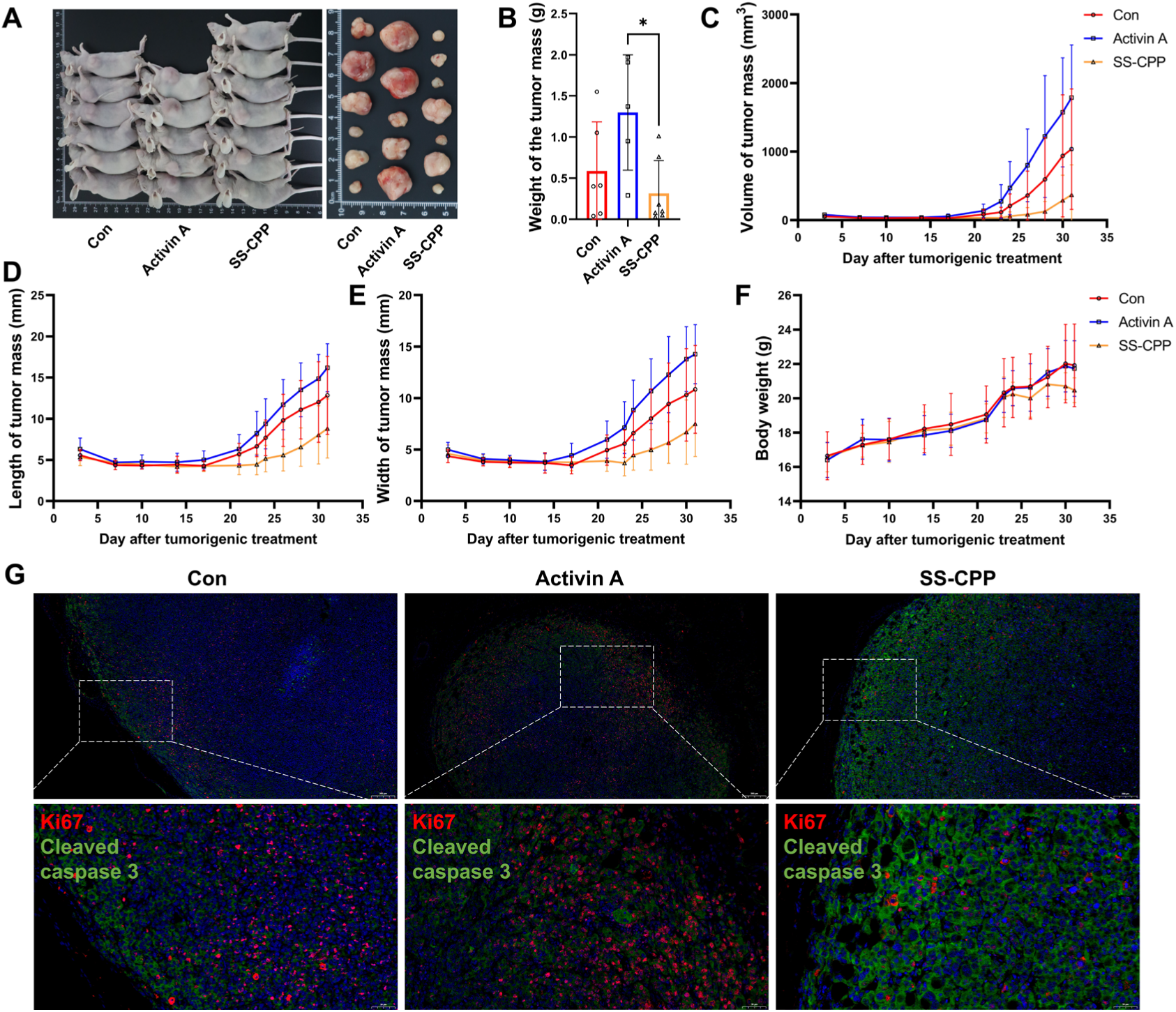
SS-CPP inhibited the growth of tumors in MCF-7 breast cancer-bearing nude mice A. Nude mice and the dissected tumor masses on the 31st day after tumor formation. Starting from the 21st day after tumor formation, the control group was injected with 0.1 mL of normal saline daily, the activin A group was injected with 0.1 mL of 100 ng/0.1 mL activin A daily, and the SS-CPP group was injected with 0.1 mL of SS-CPP synthetic polypeptide (combination of 30S-CPP and 274S-CPP) at a concentration of 75 µg/0.1 mL daily. B. Comparison of tumor weights on the 31st day after tumor formation. C-E. Growth curves of tumor masses, including the changes in tumor diameter, length, and width during the experiment. F. The change curve of the body weight of nude mice during the experiment. G. Effect of SS-CPP treatment on the proliferation and apoptosis of the formed tumor tissue. Ki67 was labeled with red fluorescence, and cleaved caspase 3 was labeled with green fluorescence. The “*” indicates a significant difference between groups.

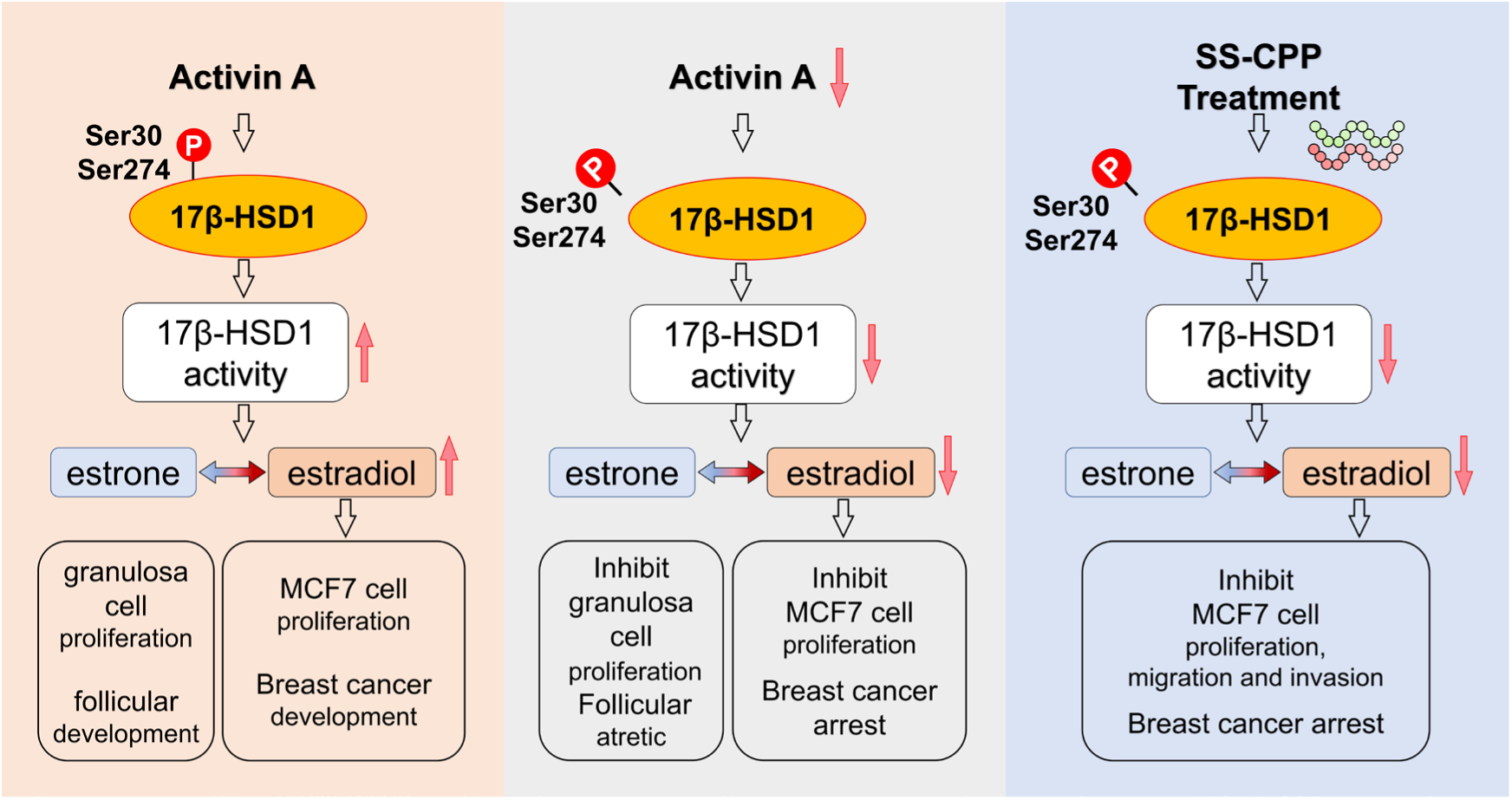

## 4 Discussion

E_2_ is an important hormone that affects the development, reproduction, life and health, and gynecological cancers. However, we have less understanding of the regulatory mechanism of the enzymatic activity of its key synthetase 17β-HSD1. This study found that phosphorylation at the key site can regulate the enzymatic activity of 17β-HSD1, and it was proved through in vitro and in vivo site mutation experiments that dephosphorylation at the key site (Ser30 and Ser274) can inhibit its enzymatic activity. Further, using follicular granulosa cells and estrogen-dependent breast cancer cell line MCF7 cells as the research objects, it was found that the follicle-stimulating factor and the oncogenic factor activin A can promote the phosphorylation of 17β-HSD1 at the key site, thereby increasing the enzymatic activity of 17β-HSD1. Then, a synthetic transmembrane peptide that can inhibit its phosphorylation and enzymatic activity was designed based on the key phosphorylation site of 17β-HSD1. Further study demonstrated that the synthetic transmembrane peptide can inhibit the phosphorylation of key sites and the enzymatic activity of 17β-HSD1, as well as suppress the proliferation and migration of granulosa cells and MCF-7 cells. Moreover, it has an inhibitory effect on tumor growth in nude mice with MCF-7 tumors. This research holds significant theoretical value and promising applications, and will greatly contribute to the advancement of animal breeding technology and human reproductive health.

The research on the enzymatic activity of 17β-HSD1 is of great significance. Cheng et., al found that ovarian stimulation-induced supraphysiological levels of E_2_ impairs uterine receptivity, thereby reducing pregnancy rate and litter size in sows ^[41]^. 17β-HSD1 was upregulated in estrogen-dependent cancers ^[3]^, therefore blocking 17β-HSD1 can be a potential novel strategy in breast cancer and endometrial cancer. In this study, two key phosphorylation site (Ser30 and Ser274) that can affect the enzymatic activity of 17β-HSD1 was innovatively discovered during the study of porcine follicular atresia ^[22]^. A new mechanism for regulating its enzymatic activity was analyzed from the perspective of post-translational modification of proteins, which has important guiding value for the development of related reproductive regulation technologies and targeted drugs.

Activin A is a multifunctional cytokine and a member of the TGF-β superfamily. The transforming growth factor-β (TGF-β) pathway plays a vital role in breast cancer metastasis ^[42]^. Activin A can stimulate granulosa cell proliferation, follicular antrum formation, and the growth and survival of follicles in vitro ^[43]^. It has been reported that activin A can regulate the enzymatic activity of 17β-HSD1, but the underline mechanism has not been fully comprehended. This research discovered that activin A and IGF-1 can upregulate the synthesis of E_2_ by enhancing the phosphorylation of the crucial sites (Ser30 and Ser274) of 17β-HSD1.

Our team discovered the key phosphorylation sites that affect 17β-HSD1 enzymatic activity during the process of follicular atresia. Then, we considered applying this finding to develop application methods for estrogen-dependent diseases. CPPs have been proven to be applicable for the delivery of various drugs. In this study, we employed CPPs to engineer synthetic transmembrane polypeptides designed to inhibit the enzymatic activity of 17β-HSD1, and investigated their functional roles in a mouse model of breast tumorigenesis. The results further demonstrated that CPPs could inhibit the enzymatic activity of 17β-HSD1 by suppressing phosphorylation at its key sites. Our findings also demonstrate that CPPs exhibit inhibitory effects on the enzymatic activity of 17β-HSD1 and suppress tumor growth in breast cancer-bearing nude mice. This further proves that phosphorylation can regulate the enzymatic activity of 17β-HSD1 and provides a new idea for the treatment of estrogen-related diseases, such as breast cancer.

This study is highly valuable as it demonstrates via a series of in vivo and in vitro experiments that phosphorylation regulates the enzymatic activity of 17β-HSD1—a finding fully validated in follicular granulosa cells and breast cancer cells. The synthetic cell-penetrating peptides (especially SS-CPP) designed in this study have the function of inhibiting the enzymatic activity of 17β-HSD1 and reducing the synthesis of estradiol, which is of great guiding value for the development of drugs for cancers such as breast cancer. In this study, linear synthetic cell-penetrating peptides were used. In future pharmaceutical research and development, cyclic synthetic cell-penetrating peptides can be considered to improve the half-life and practicality of drugs, as cyclic peptides have increased proteolytic stability and bioavailability ^[44, 45]^. In addition, a limitation of this study is that point mutations and enzymatic activity studies were directly conducted in eukaryotic cells without prokaryotic expression of the protein.

## 5 Conclusion

Our study found that through in vitro site-directed mutagenesis and experiments on point-mutated mice, Ser30 and Ser274 were identified as key phosphorylation sites that significantly affect the enzymatic activity of 17β-HSD1. We demonstrated that activin A and IGF-1, factors that promote E_2_ synthesis, enhance the enzymatic activity of 17β-HSD1 by increasing its phosphorylation at these key sites. Based on these key phosphorylation sites of 17β-HSD1, the synthetic cell-penetrating peptides we designed (particularly SS-CPP) inhibit the enzymatic activity of 17β-HSD1 by suppressing its phosphorylation and also inhibit tumor growth in a mouse breast cancer model. The findings of this study offer new insights into the development of novel reproductive technologies in mammals and the treatment of estrogen-dependent diseases, including breast cancer.

## Data availability statement

The proteome data for porcine GCs from healthy, slightly atretic, and atretic follicles were obtained from DOI: https://doi.org/10.3389/fcell.2020.624985. The mass spectrometry proteomics data have been deposited to the ProteomeXchange Consortium via the PRIDE partner repository with the dataset identifier PXD020899. The raw data of “The Impact of SS-CPP Treatment on the Gene Expression Profile of MCF-7 Cells” have been deposited in the Sequence Read Archive (SRA) database with BioProject accession number PRJNA1345717. All other raw data are available from the corresponding author upon reasonable request.

## Acknowledgment

We would like to express our gratitude to the laboratory of Dr. Yongpeng Guo at the College of Animal Science and Technology, Henan Agricultural University, for providing the solution for detecting estradiol by high-performance liquid chromatography. We also acknowledge the College of Veterinary Medicine, Henan Agricultural University, for providing the animal housing facility.

## Author contributions

H.F., S. C., X. L., Y. L., K. L., and H. W. conducted the experiments, S. Z., K. W., X. H., J. L., X. L., and F. Y. designed the experiments. Funding acquisition: J. L. and F. Y.. Supervision: X. L., and F. Y.. Writing–original draft: H.F., S. C., and F. Y.. Writing–review & editing: F. Y..

## Ethics approval

All trials and investigations were approved by the Ethical Committee of Henan Agricultural University (Ethical approval number: HNND2022030856).

