## Supplementary Figures for "A novel dephosphorylation peptide inhibits 17β-HSD1 enzyme activity in ovarian granulosa cells and breast cancer cells"

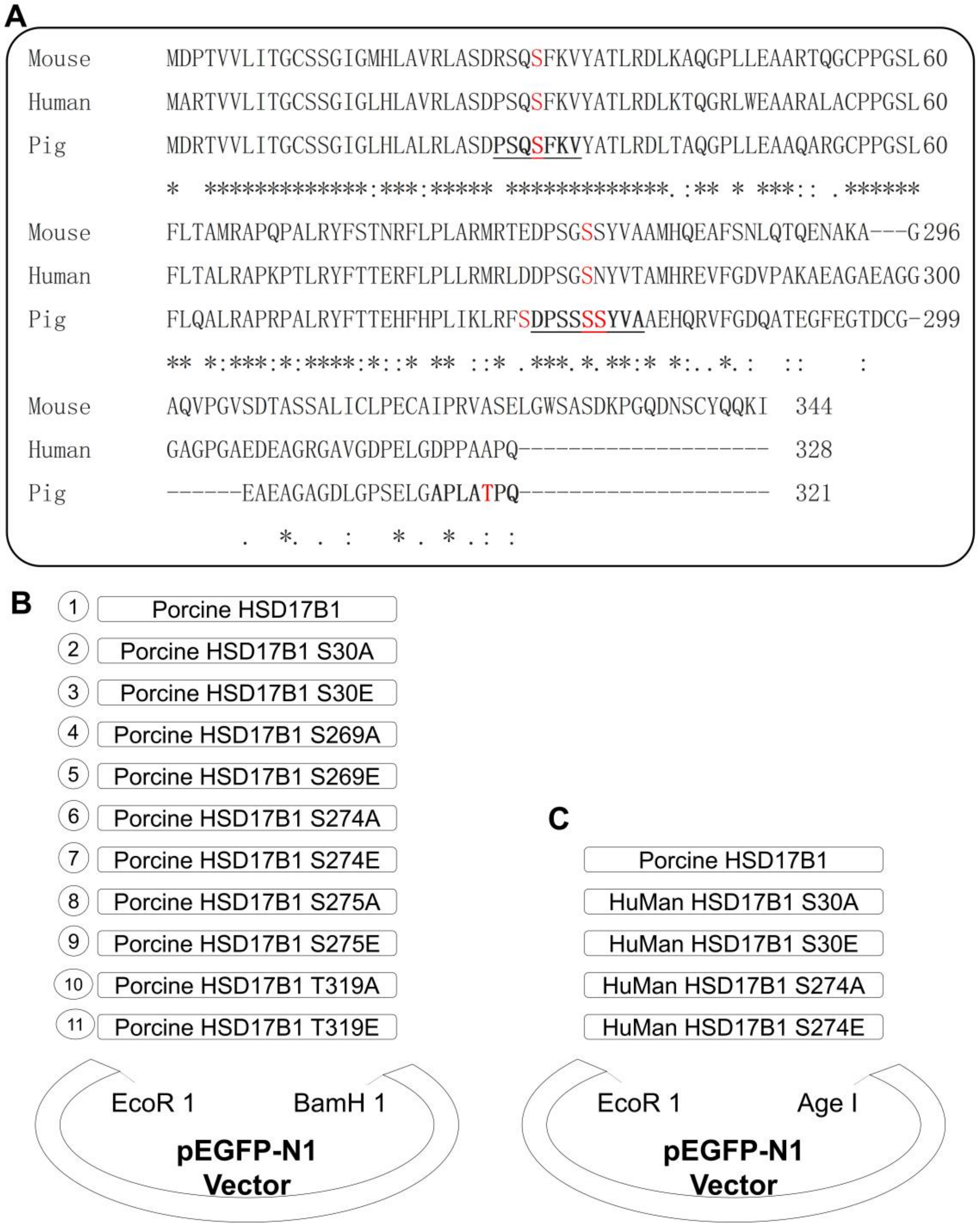


Fig. S1. Inter-species alignment of 17β-HSD1 amino acid sequences and construction of HSD17B1 expression vectors

A. Alignment of 17β-HSD1 amino acid sequences among three species: human, pig, and mouse. B. Schematic diagram of the construction of porcine HSD17B1 and their key site mutant vectors. C. human HSD17B1 and their key site mutant vectors. The figure shows information such as the names of the vectors and restriction enzymes used.

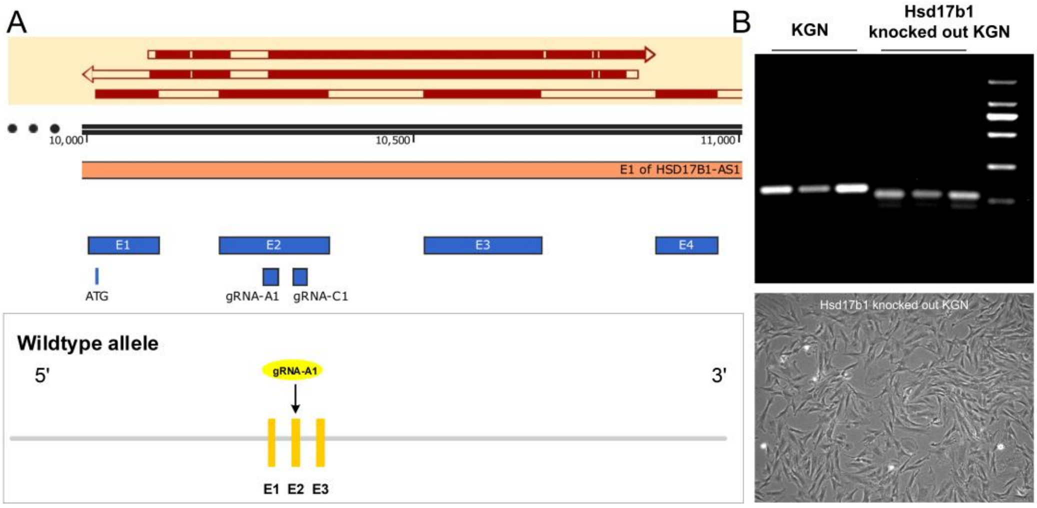


Fig. S2. Construction of Hsd17b1 knocked out KGN cells and verification of their genotypes

A. The Human HSD17B1 gene in ovarian granulosa cell line KGN was knocked out by CRISPR/Cas9 technology. gRNA-A1: ACGTCTCCAGGGATCCCGGA - GGG. After electroporation, single clones were picked and verified by PCR and sequencing, and homozygous cells with the Human HSD17B1 gene knockout were successfully obtained. B. Genotype identification. PCR and sequencing primers: upstream primer (F1): TTGGCCGTACGTCTGGCTT, downstream primer (R1): TTAGGTGGGGAGACGAGGTT. The sequencing results showed a 58 - bp base deletion, which was consistent with the DNA gel electrophoresis results.

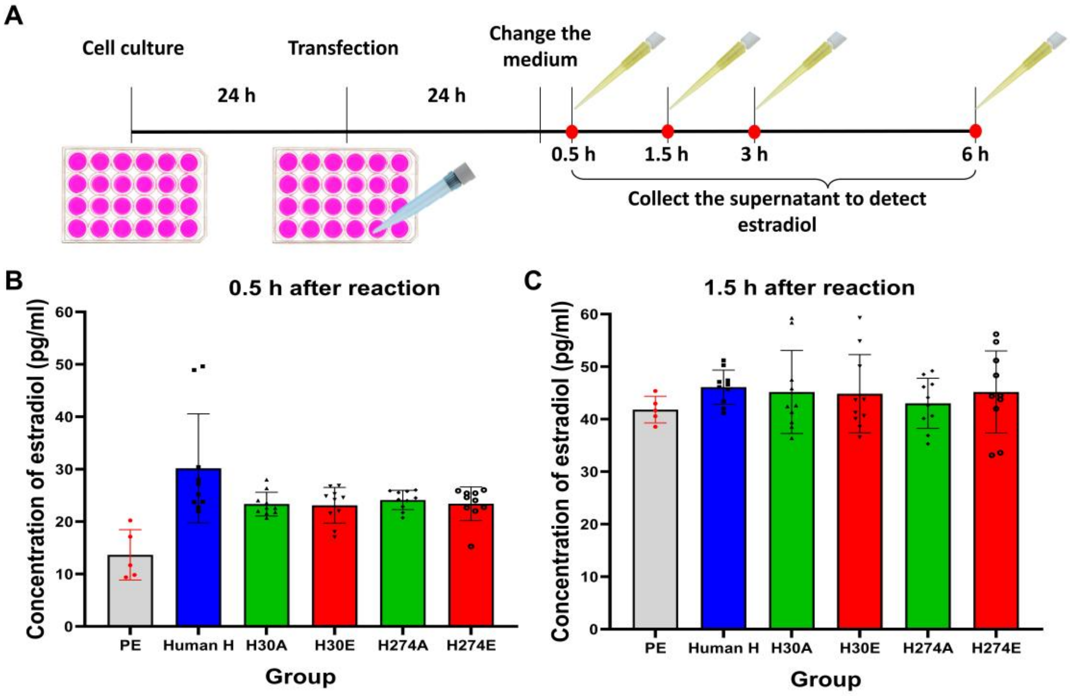

Fig. S3. Effect of transfecting human 17β-HSD1 and its mutant vectors into 293T cells not significantly influence $E_2$ conversion rate

A. Flow chart of the experiment of transfecting human 17β-HSD1 and its mutant vectors into 293T cells and detecting $E_2$ synthesis. After transfection, 1 ng/mL $E_1$ was added as a substrate to the cell culture medium, and $E_2$ production was measured at different time points after addition. B and C, The amount of $E_2$ synthesized after 0.5 and 1.5 hours of reaction with 1 ng/mL $E_1$.

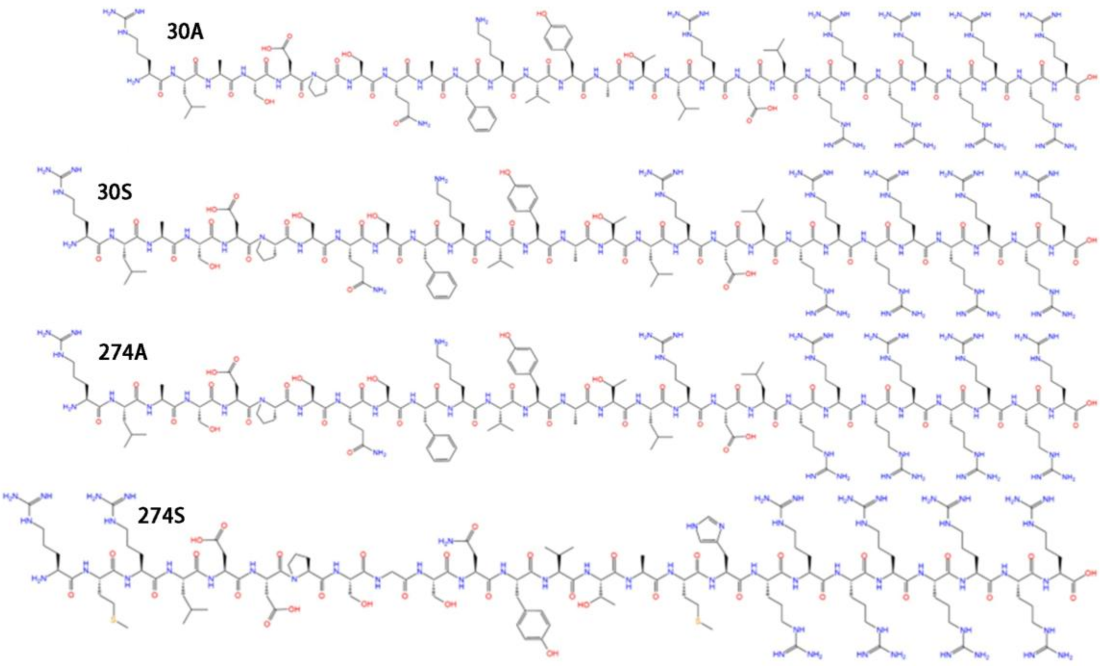


**Fig. S4**. The molecular structural formula of the transmembrane polypeptide

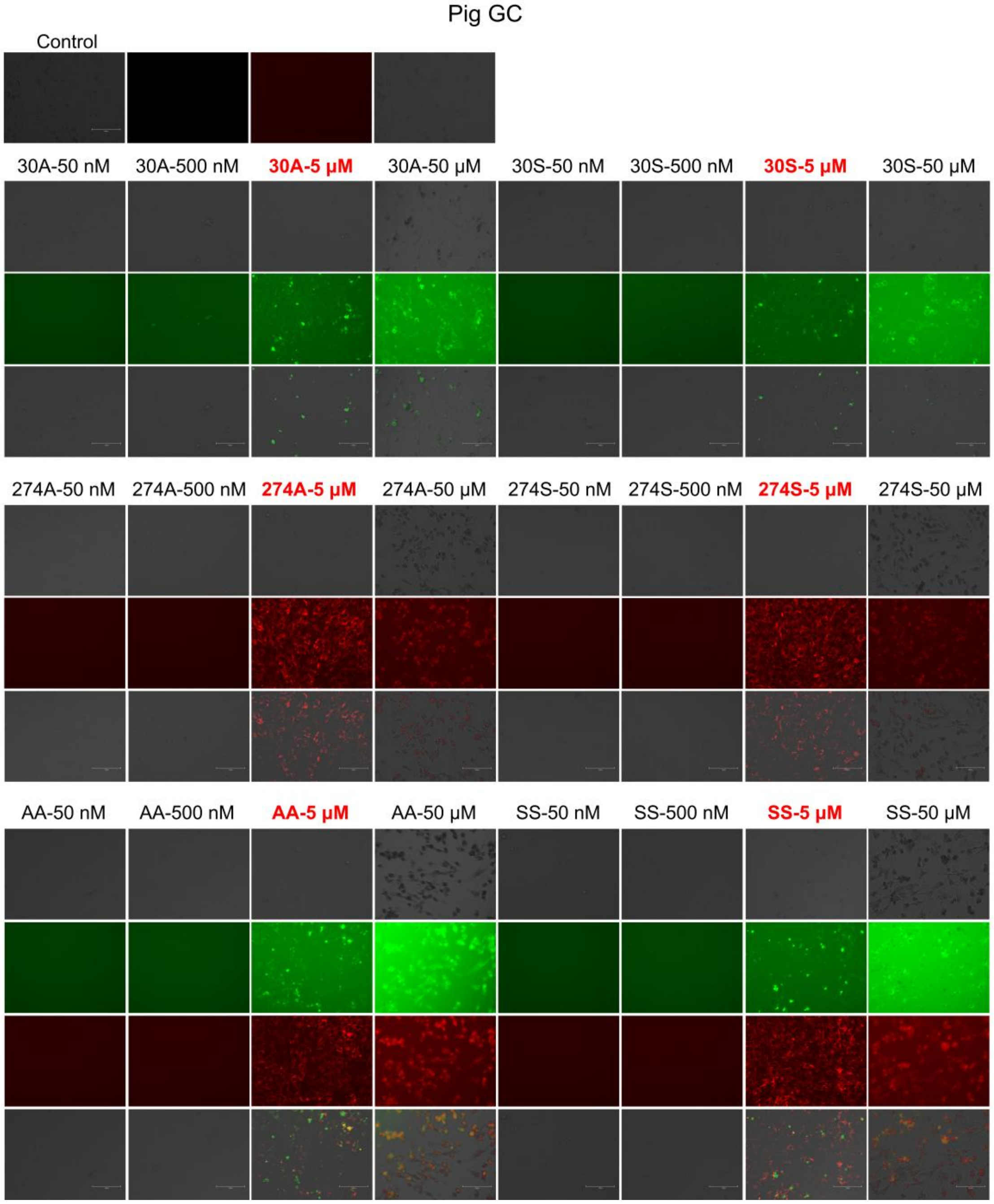


**Fig. S5.** The cell-penetrating ability of transmembrane polypeptides at different concentrations in the pig granulosa cells.

30A= S30A-CPP, 30S= S30-CPP, 274A= S274A-CPP, 274A-CPP= S274-CPP, AA=combined use of S30A-CPP and S274A-CPP, SS= combined use of S30-CPP and S274-CPP.

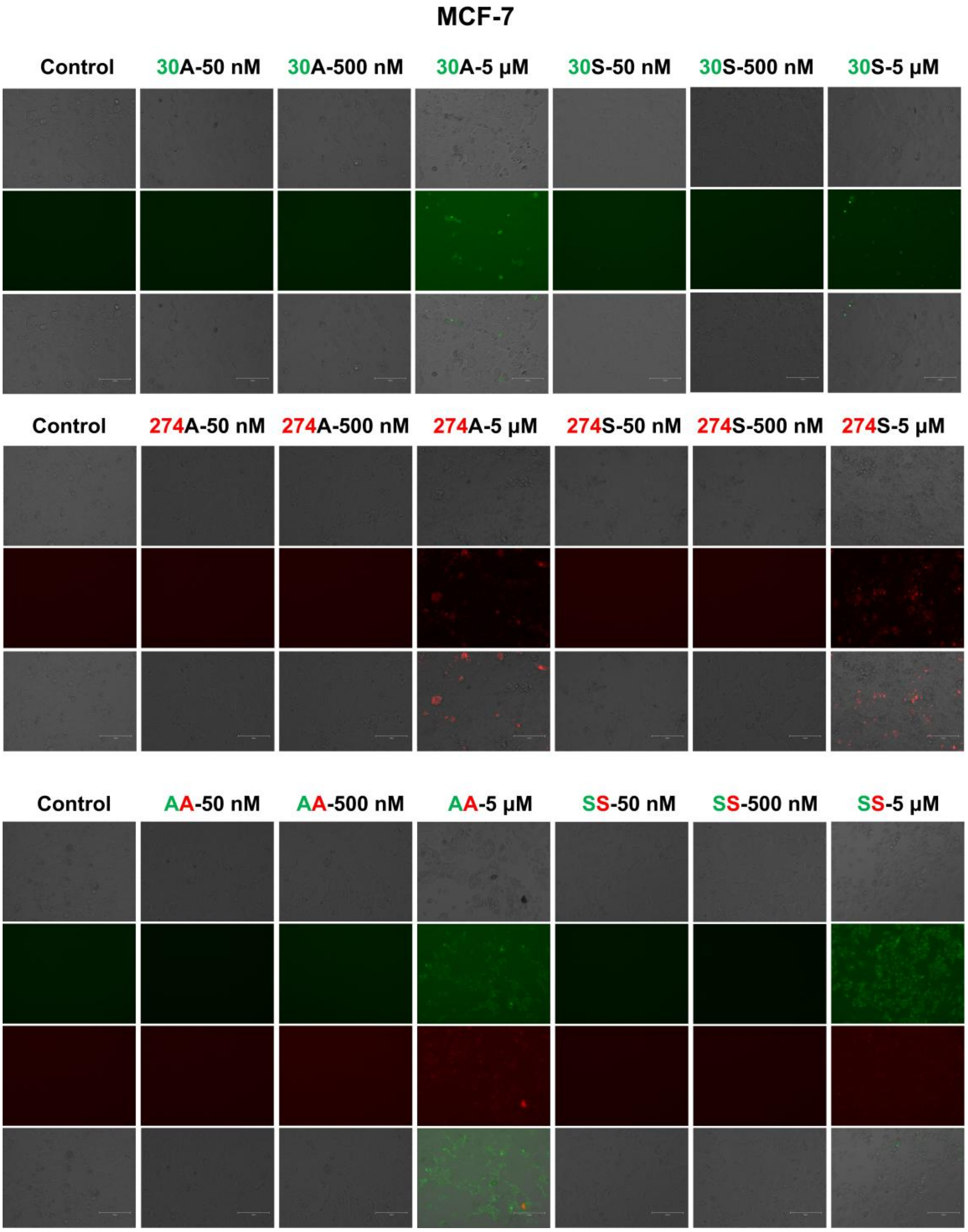


**Fig. S6.** The cell-penetrating ability of transmembrane polypeptides at different concentrations in the MCF-7 cells.

30A= S30A-CPP, 30S= S30-CPP, 274A= S274A-CPP, 274A-CPP= S274-CPP, AA=combined use of S30A-CPP and S274A-CPP, SS= combined use of S30-CPP and S274-CPP.
