## Supplementary Table 1 for "A novel dephosphorylation peptide inhibits 17β-HSD1 enzyme activity in ovarian granulosa cells and breast cancer cells"

**Supplementary Table 1.** The primers for amplifying HSD17B1 and performing point mutations on it.

| | Primer Name | Primer Sequence |
|---|---|---|
| pig HSD17B1 | PE ECOR1 pig HSD17B1 forward | GAATTCatggaccgcaccgtggtgctcat |
| | PE BamH1 pig HSD17B1 reverse | GGATCCtattgcggggtggcaaga |
| | pig HSD17B1 S30A forward | tctgacccatctcagGCAttcaaagtgtacg |
| | pig HSD17B1 S30A reverse | TGCctgagatgggtcagatgccagacgcag |
| | pig HSD17B1 S30E forward | tctgacccatctcagGAGttcaaagtgtacg |
| | pig HSD17B1 S30E reverse | CTCctgagatgggtcagatgccagacgcag |
| | pig HSD17B1 S274A forward | tccgaccccagcagcGCAagctacgtcgcagcc |
| | pig HSD17B1 S274A reverse | TGCgctgctggggtcggagaagcgcagcttga |
| | pig HSD17B1 S274E forward | tccgaccccagcagcGAGagctacgtcgcagcc |
| | pig HSD17B1 S274E reverse | CTCgctgctggggtcggagaagcgcagcttga |
| | pig HSD17B1 S275A forward | gaccccagcagcagcGCAtacgtcgcagccga |
| | pig HSD17B1 S275A reverse | TGCgctgctgctggggtcggagaagcgcagctt |
| | pig HSD17B1 S275E forward | gaccccagcagcagcGAGtacgtcgcagccga |
| | pig HSD17B1 S275E reverse | CTCgctgctgctggggtcggagaagcgcagctt |
| | pig HSD17B1 T319A forward | GAATTCatggaccgcaccgtggtgctcat |
| | pig HSD17B1 T319A reverse | GGATCCtattgTgcggtggcaagag |
| | pig HSD17B1 T319E forward | GAATTCatggaccgcaccgtggtgctcat |
| | pig HSD17B1 T319E reverse | GGATCCtattgcTCggtggcaagag |
| human HSD17B1 | Human HSD17B1 forward 1 | GAATTCATGGCCCGCACCGTGGTGCTCAT |
| | Human HSD17B1 reverse 1 | ACCGGTgcCTGCGGGGCGGCCGGAGGATCGCCGAGCTCA |
| | Human HSD17B1 S30A forward | TCAGATCCATCCCAGGCCTTCAAAGTGTATGC |
| | Human HSD17B1 S30A reverse | GGCCTGGGATGGATCTGAAGCCAGACGTAC |
| | Human HSD17B1 S30E forward | TCAGATCCATCCCAGGAGTTCAAAGTGTATGC |
| | Human HSD17B1 S30E reverse | CTCCTGGGATGGATCTGAAGCCAGACGTAC |
| | Human HSD17B1 S274A forward | TGGACGACCCCAGCGGCGCCAACTACGTCACCGCCA |
| | Human HSD17B1 S274A reverse | GGCGCCGCTGGGGTCGTCCAGGCGCATCCGCAGCAG |
| | Human HSD17B1 S274E forward | TGGACGACCCCAGCGGCGAGAACTACGTCACCGCCA |
| | Human HSD17B1 S274E reverse | CTCGCCGCTGGGGTCGTCCAGGCGCATCCGCAGCAG |
